# Underwater CAM photosynthesis elucidated by *Isoetes* genome

**DOI:** 10.1101/2021.06.09.447806

**Authors:** David Wickell, Li-Yaung Kuo, Hsiao-Pei Yang, Amra Dhabalia Ashok, Iker Irisarri, Armin Dadras, Sophie de Vries, Jan de Vries, Yao-Moan Huang, Zheng Li, Michael S. Barker, Nolan T. Hartwick, Todd P. Michael, Fay-Wei Li

**Affiliations:** Plant Biology Section, School of Integrative Plant Science, Cornell University, Ithaca, NY, USA; Boyce Thompson Institute, Ithaca, NY, USA; Institute of Molecular & Cellular Biology, National Tsing Hua University, Hsinchu, Taiwan; Institute for Microbiology and Genetics, Department of Applied Bioinformatics, University of Goettingen, Goettingen, Germany; Campus Institute Data Science, University of Goettingen, Goettingen, Germany; Goettingen Center for Molecular Biosciences, Department of Applied Bioinformatics, University of Goettingen, Goettingen, Germany; Taiwan Forestry Research Institute, Taipei, Taiwan; Department of Integrative Biology, University of Texas at Austin, Austin, TX, USA; Department of Ecology and Evolutionary Biology, University of Arizona, Tucson, AZ, USA; The Molecular and Cellular Biology Laboratory, The Salk Institute for Biological Studies, La Jolla, CA, USA

## Abstract

To conserve water in arid environments, numerous plant lineages have independently evolved Crassulacean Acid Metabolism (CAM). Interestingly, *Isoetes*, an aquatic lycophyte, can also perform CAM as an adaptation to low CO_2_ availability underwater. However, little is known about the evolution of CAM in aquatic plants and the lack of genomic data has hindered comparison between aquatic and terrestrial CAM. Here, we investigated the underwater CAM in *Isoetes taiwanensis* by generating a high-quality genome assembly and RNA-seq time course. Despite broad similarities between CAM in *Isoetes* and terrestrial angiosperms, we identified several key differences. Notably, for carboxylation of PEP, *Isoetes* recruited the lesser-known “bacterial-type” PEPC, along with the “plant-type” exclusively used in other terrestrial CAM and C4 plants. Furthermore, we found that circadian control of key CAM pathway genes has diverged considerably in *Isoetes* relative to flowering plants. This suggests the existence of more evolutionary paths to CAM than previously recognized.

---

*Isoetes*, commonly known as quillworts, is the only genus in the lycophyte order Isoetales, containing roughly 250 described species^1^. It is the last remaining member of an ancient lineage with a fossil record that dates back to at least the late Devonian. As such, quillworts are believed to represent the closest living relatives of the giant, tree-like lycopsids such as *Sigillaria* and *Lepidodendron* that dominated the terrestrial landscape during the Carboniferous^2^. However, in contrast to its arborescent ancestors, modern *Isoetes* species are diminutive and mostly aquatic with the vast majority of species growing completely or partially submerged. Underwater, *Isoetes* can conduct CAM^3^, a carbon concentrating mechanism involving the separation of carbon uptake and fixation in a time of day (TOD) fashion, with carbon being sequestered as malate at night, to be fed into the Calvin cycle during the day. CAM is a common strategy to improve water-use efficiency among xeric-adapted plants, allowing them to keep their stomata closed during the day. However, its prevalence in aquatic species of *Isoetes*^3^, as well as several aquatic angiosperms^4,5^, suggests that it must have some utility unrelated to conserving water. Specifically, it is thought to be an adaptation to low aquatic CO_2_ availability in the oligotrophic lakes and seasonal pools where *Isoetes* species are commonly found^4,6^.

Though it has been nearly four decades since Keeley first described “CAM-like diurnal acid metabolism” in *Isoetes howellii*^7^, relatively little is known about the genetic mechanisms controlling CAM in *Isoetes* or any other aquatic plant. Previous genomic and/or transcriptomic studies that focused on terrestrial CAM have found evidence for regulatory neofunctionalization, enrichment of cis-regulatory elements, and/or reprogramming of gene regulatory networks that underlie the convergent evolution of CAM in *Sedum album*^8^, *Ananas comosus*^9^, *Kalanchoe fedtschenkoi*^10^, several orchids^11–13^, and Agavoideae species^14,15^. Furthermore, a remarkable case of amino acid sequence convergence in phosphoenolpyruvate carboxylase (PEPC), which catalyzes the carboxylation of phosphoenolpyruvate (PEP) to yield oxaloacetate (OAA), has also been reported among terrestrial CAM plants^10^. However, the lack of a high-quality genome assembly has made meaningful comparison of *Isoetes* or any other aquatic CAM plant to terrestrial CAM species impossible.

The only lycophyte genomes available to date are from the genus *Selaginella*^16–18^, leaving a deep, >300-million-year gap in our knowledge of lycophyte genomics and limiting inferences of tracheophyte evolution. *Selaginella* is the only genus in the Selaginellales, the sister clade to Isoetales. Notably, *Selaginella* is known for being one of few lineages of vascular plants for which no ancient whole genome duplications (WGDs) have been detected. Conversely, there is evidence from transcriptomic data for as many as two rounds of WGD in *Isoetes tegetiformans*^19^. As such, a thorough characterization of the history of WGD in *Isoetes* is vital to future research into the effects and significance of WGD in lycophytes as a whole.

With this study we sought to investigate genome evolution as well as the genetic underpinnings of CAM in *Isoetes*. To that end, we present a high-quality genome assembly for *Isoetes taiwanensis*. To our knowledge, this is not only the first such assembly for the order Isoetales, but also the first for an aquatic CAM plant. We found evidence for a single ancient WGD event that appears to be shared among multiple species of *Isoetes*. Additionally, while many CAM pathway genes display similar expression patterns in *Isoetes* and terrestrial angiosperms, notable differences in gene expression suggest that the evolution of CAM may have followed very different trajectories in these highly divergent groups.

## Results and Discussion

### Genome assembly, annotation, and organization

Using Illumina short-reads, Nanopore long-reads, and Bionano optical mapping, 90.13% of the diploid (2n = 2x = 22 chromosomes) *I. taiwanensis* genome was assembled into 204 scaffolds (N50=17.40 Mb), with the remaining 9.87% into 909 unplaced contigs (Table 1). The total assembled genome size (1.66 Gb) is congruent with what was estimated by K-mers (1.65 Gb) and flow cytometry (1.55 Gb) (Supplementary Fig. S1). A circular-mapping plastome was also assembled, from which we identified a high level of RNA-editing (Supplementary Notes, Supplementary Fig. S2).

**Table 1.** *Isoetes taiwanensis* genome assembly statistics.

|  |  |
| --- | --- |
| Assembly size (Mb) | 1,658.30 |
| Scaffolds (#) | 204 |
| Scaffold length (Mb) | 1494.58 |
| N50 of scaffold length (Mb) | 17.40 |
| Scaffolded contigs (#) | 1,879 |
| Scaffolded contig length (Mb) | 1211.25 |
| N50 length of scaffolded contigs (Mb) | 1.48 |
| Un scaffolded contig (#) | 909 |
| Unscaffolded contig (Mb) | 149.46 |
| N50 length of unscaffolded contigs (Mb) | 0.26 |
| Genome BUSCO score (Eukaryota) (%) | 94.5 |
| Proteome BUSCO score (Eukaryota) (%) | 91.0 |
| Predicted protein coding genes (#) | 39,461 |
| Predicted repetitive sequence (%) | 38 |

A total of 39,461 high confidence genes were annotated based on *ab initio* prediction, protein homology, and transcript evidence. The genome and proteome BUSCO scores are 94.5% and 91.0% respectively, which are comparable to many other seed-free plant genomes (Supplementary Fig. S3) and indicative of high completeness. Orthofinder^20^ analysis of 25 genomes placed 647,535 genes into 40,144 orthogroups. Subsequent analysis of key stomatal and root genes in *I. taiwanensis* genome supported their homology (at the molecular level) with similar structures in other vascular plants. In addition, examination of lignin biosynthesis genes in *I. taiwanensis* suggests that evolution of a novel pathway to S-lignin likely predates the divergence of *Isoetes* and *Selaginella*. A detailed discussion of these analyses and Orthofinder results can be found in the Supplementary Notes and Supplementary Figures S4-S20.

Repetitive sequences accounted for 38% of the genome assembly with transposable elements (TEs) accounting for the majority of those at 37.08% of the assembly length. Long terminal repeat (LTR) retrotransposons were the most abundant (15.72% of total genome assembly) with the Gypsy superfamily accounting for around 68% of LTR coverage (10.7% of total genome assembly; Supplementary Table 1). When repeat density was plotted alongside gene density, the distribution of both was found to be homogeneous throughout the assembly (Fig. 1). This even distribution of genes and repeats is markedly different from what has been reported in most angiosperm genomes^21^ where gene density increases near the ends of individual chromosomes, but consistent with several high-quality genomes published from seed-free plants, including *Physcomitrium patens*^22^, *Marchantia polymorpha*^23^, and *Anthoceros agrestis*^24^. The result from *I. taiwanensis* thus adds to the growing evidence that the genomic organization might be quite different between seed and seed-free plants^25^.

**Fig 1:**
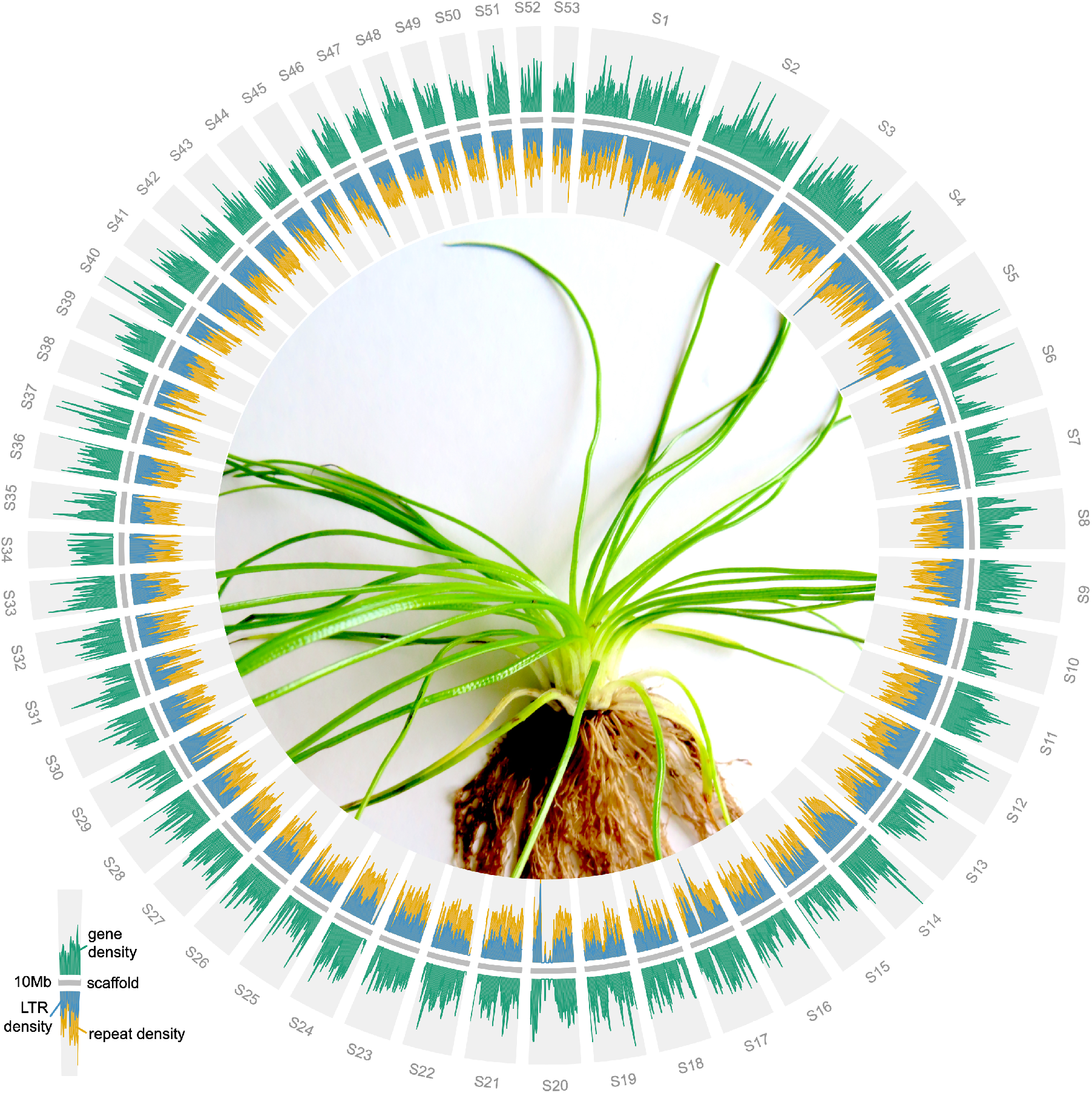
Distribution of genes and repetitive elements in *I. taiwanensis*. The relatively even distributions differ from angiosperm genomes, but are similar to what have been reported in other seed-free plants. Only scaffolds longer than 10 Mb are plotted. Center: an image of *I. taiwanensis*.

### Evidence for WGD in Isoetes taiwanensis

Using a combination of methods including synonymous substitutions per site (Ks), phylogenetic, and synteny analyses, we identified a single ancient WGD in *I. taiwanensis*. This is in contrast to a previous Ks analysis using 1KP transcriptome data, which found evidence for two rounds of WGD, named ISTEα and ISTEβ, in the North American species *I. tegetiformans* and *I. echinospora*^26^. These two WGDs have median Ks values of ∼0.5 and ∼1.5^26^ (Supplementary Figure S21). Our whole paranome analysis of Ks in *I. taiwanensis* revealed a single peak at Ks ∼ 1.8 (Fig. 1a), suggesting that the earlier of the two duplications (ISTEβ) in *I. tegetiformans* and *I. echinospora* is shared by *I. taiwanensis* while the more recent event (ISTEα) is not. Further analysis of orthologous divergence between *I. taiwanensis* and *I. lacustris* indicated that ISTEβ predates the divergence of these two species (Supplementary Figure S22). The ISTEβ event was subsequently confirmed by gene tree-species tree reconciliation using genomic data in the WhALE package^27^. WhALE returned a posterior distribution of gene retention centered on q = ∼0.12. This result compares favorably with a previously documented WGD event in *Azolla filiculoides*^28^ (q = ∼0.08) and is in stark contrast to our negative control, *Marchantia polymorpha*^*23*^ (q = ∼0) (Fig. 2b,c).

**Fig. 2:**
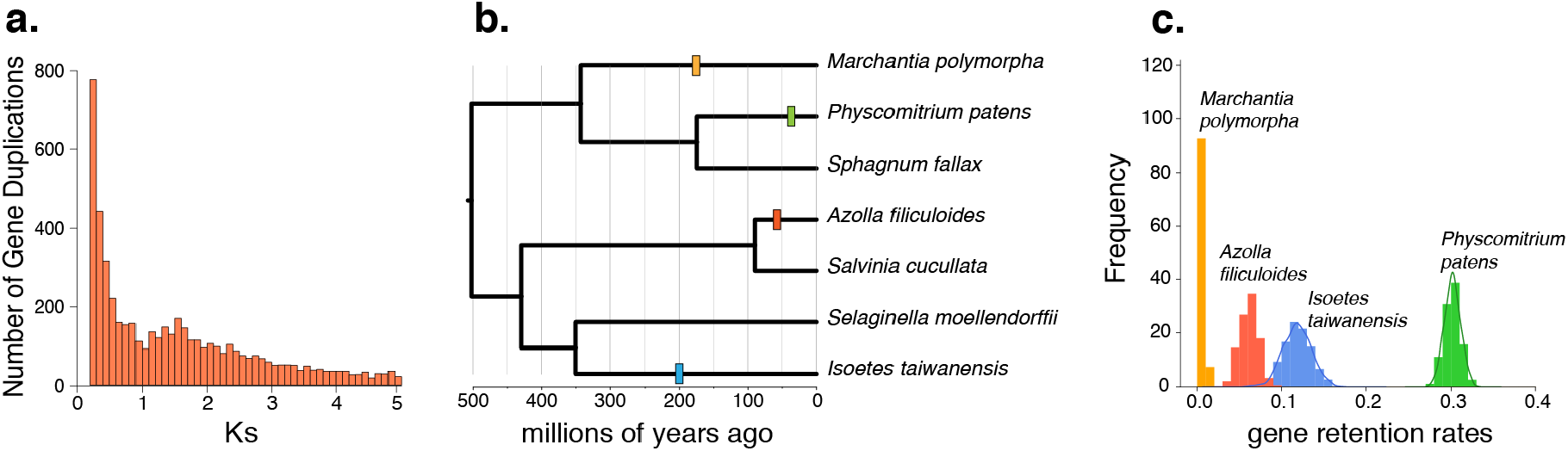
Evidence for WGD in *I. taiwanensis*. **a**, Ks plot showing a peak centered on 1.8 corresponding to the ISTEβ event. **b**, Hypothesized WGD events that were tested (colored rectangles) in our WhALE analysis are shown on a phylogeny. **c**, *I. taiwanensis’* posterior distribution of gene retention rates falls between that of *A. filiculoides* and *P. patens*, both are known to have at least one WGD. This provides additional support for the ISTEβ event. Conversely, the gene retention rate is close to zero for *M. polymorpha*, consistent with its lack of WGD.

While self-self syntenic analysis revealed 6,196 genes (15.7%) with a syntenic depth of 1x in 107 clusters (Supplementary Figure S23), we do not believe they resulted from WGD. Our Ks analysis restricted to syntenic gene pairs failed to recover the peak at Ks ∼1.8 and instead consisted of an initial slope toward a much lower Ks value (Supplementary Figure S24). Considering their high degree of similarity and location on separate scaffolds, it is possible that these low Ks gene pairs are the result of relatively recent segmental duplications. The absence of synteny from ISTEβ is unsurprising. The high Ks value implies that ISTEβ is quite ancient; long enough ago for extensive genomic restructuring and fractionation to have taken place. Altogether, of the two hypothesized WGDs in *Isoetes*, we confirmed the presence of ISTEβ while the younger ISTEα might be either specific to *I. tegetiformans* and *I. echinospora* or an artifact stemming from the quality or completeness of the transcriptomes.

### Similarities to terrestrial CAM plants

As a lycophyte, *Isoetes* represents the oldest extant lineage of vascular plants to exhibit CAM photosynthesis (Fig 3a), and may be considered unusual among other CAM plants due to its aquatic lifestyle. Here, we demonstrated that when submerged, titratable acidity in the leaves of *I. taiwanensis* increased throughout the night, reaching peak acidity in the morning and decreased throughout the daylight hours (Fig 3b), consistent with the cycle of carbon sequestration and assimilation seen in dry-adapted CAM plants. To identify the underlying genetic elements, we generated TOD RNA-seq, sampling every 3 hours over a 27-hour period under 12 h light/12 h dark and continuous temperature (LDHH). A multidimensional scaling (MDS) plot of normalized expression data showed that the samples were generally clustered in a clockwise fashion as expected for TOD expression analysis (Supplemental Figure S25).

**Fig. 3:**
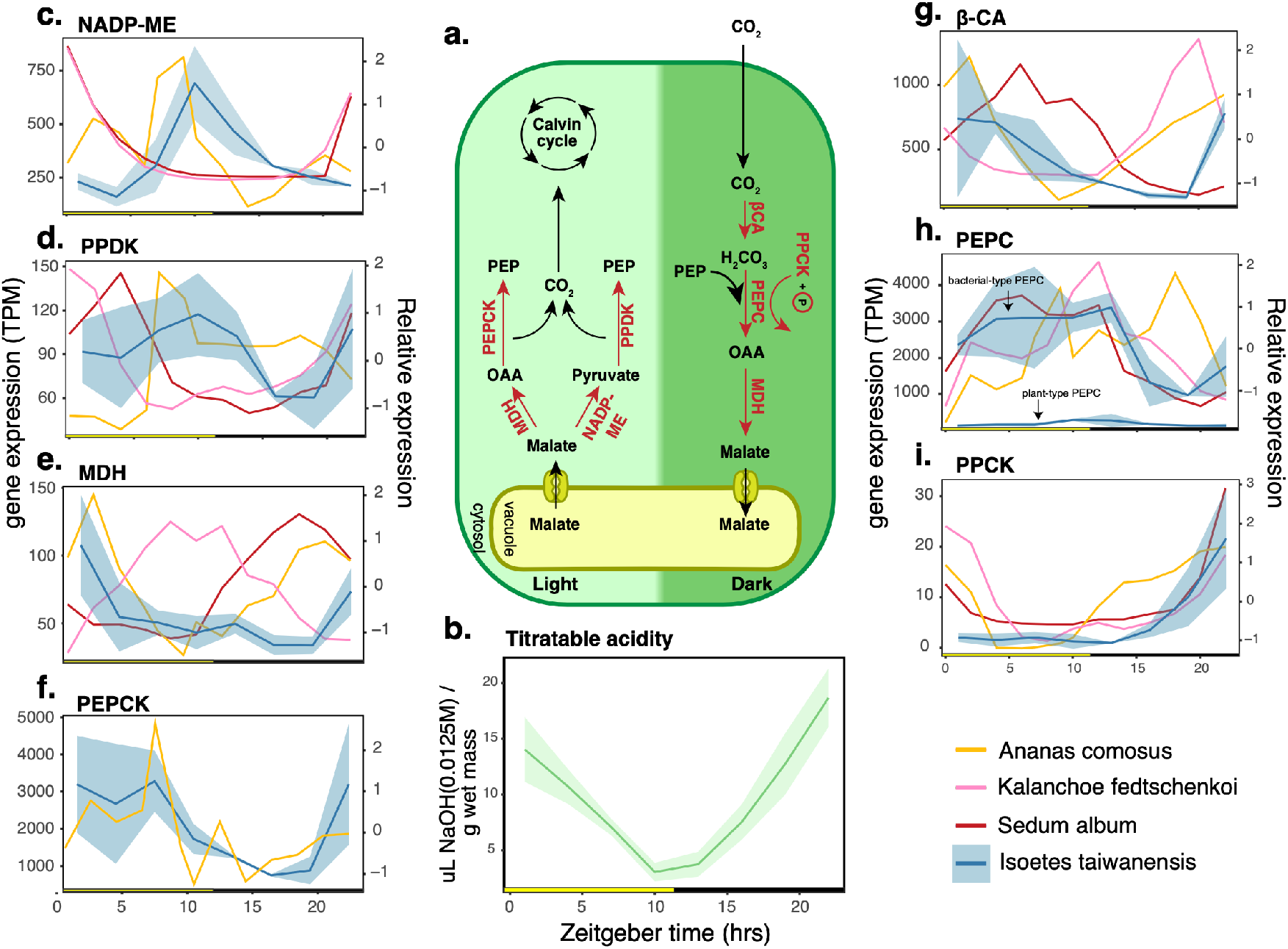
Key CAM pathway genes and their expression patterns in *I. taiwanensis*. **a**, The CAM pathway with important reactions and their enzymes shown in red. **b**, Titratable acidity in *I. taiwanensis* exhibited a clear diel fluctuation. Diel expression patterns for highlighted genes are shown for the day (**c-f**) and night reactions (**g-i**). Average of TPM normalized expression data for *I. taiwanensis* is plotted in blue with a shaded ribbon representing the standard deviation (left y-axis). Relative expression profiles for homologous, cycling genes in other CAM species are plotted for comparison (right y-axis). All times are displayed in hours after lights-on (Zeitgeber time).

We found that some of the CAM pathway genes in *I. taiwanensis* exhibited TOD expression patterns that largely resemble those found in terrestrial CAM plants (Fig. 3c-i). For example, the strong dark expression of *PHOSPHOENOLPYRUVATE CARBOXYLASE KINASE* (*PPCK*) appears to be conserved in *I. taiwanensis* as well as in all three terrestrial taxa (Fig 3i). Likewise, we found one copy of *β-CARBONIC ANHYDRASE* (*β-CA*) that cycled similarly with homologs in *A. comosus* and *K. fedtschenkoi* (Fig. 3g)—increasing during the night and peaking in early morning—although this is different from *S. album* in which no *β-CA* genes showed a high dark expression. Similar to *A. comosus* where two copies of *MALATE DEHYDROGENASE* (*MDH*) were found to cycle in green leaf tissue^9^, we found multiple copies of *MDH* that appear to cycle in *I. taiwanensis* with one copy appearing to exhibit similar peak expression to its orthologue in pineapple (Fig. 3e). However, neither of the other two *MDH* genes that cycle in *I. taiwanensis* exhibit similar expression to their orthologues in terrestrial CAM species (Supplementary Figure S26).

During the day, decarboxylation typically occurs by one of two separate pathways (Fig. 3a). The first utilizes *NAPD-MALIC ENZYME* (*NADP-ME*) and *PYRUVATE PHOSPHATE DIKINASE* (*PPDK*), and appears to be favored by *K. fedtschenkoi* and *S. album*^8,10^. The second utilizes *MDH* and *PHOSPHOENOLPYRUVATE CARBOXYKINASE* (*PEPCK*) and is favored by *A. comosus*^9^. Based on its TOD expression of multiple copies of *MDH* and associated expression dynamics, it is possible that *I. taiwanensis* utilizes the *MDH*/*PEPCK* pathway. While all four genes have elevated expression levels during the day, the expression of *NADP-ME* is inverted compared to *K. fedtschenkoi* and *S. album* (Fig. 3c), and *PPDK* exhibits relatively weak cycling overall (R=0.637; Fig. 3d). Additionally, *PEPCK* and one copy of *MDH* have similar TOD expression in *I. taiwanensis* and *A. comosus* (Fig. 3f and 3e, respectively), which may indicate a shared affinity for *MDH*/*PEPCK* decarboxylation. Interestingly, the copy of *PEPCK* that cycles in *I. taiwanensis* is not orthologous to the copy that cycles in *A. comosus*, being placed in a different orthogroup by Orthofinder^20^.

### I. taiwanensis has recruited bacterial-type PEPC

While TOD expression of many key CAM pathway genes was broadly similar to that seen in terrestrial CAM plants, one important difference can be found in the PEPC enzyme, which is the entry point of carboxylation in CAM and C4 photosynthesis (Fig. 3a). PEPC is present in all photosynthetic organisms as well as many non-photosynthetic bacteria and archaea. It is a vital component of plant metabolism, carboxylating PEP in the presence of HCO_3_^-^ to yield OAA. In plants, the *PEPC* gene family consists of two clades, the “plant-type” and the “bacterial-type.” The latter was named because of its higher sequence similarity with proteobacteria *PEPC* than other plant-type *PEPC* genes^29^. All CAM and C4 plants characterized to date recruited only the plant-type *PEPC*^30^, with the bacterial-type often being expressed at relatively low levels and/or primarily in non-photosynthetic tissues^31^.

Interestingly, in *I. taiwanensis* we found that both types of *PEPC* were cycling and that the bacterial-type was expressed at much higher levels than plant-type *PEPC* (Fig. 3h). Copies from both types had similar expression profiles in *I. taiwanensis*, peaking at dusk and gradually tapering off during the night. While this may seem counterintuitive as PEPC is an important component of the dark reactions, it is consistent with what has previously been found in other terrestrial CAM plants, with the overall expression profile resembling that of *S. album*^8^. The advantage of recruiting bacterial-type *PEPC* is unclear. *In vivo*, both bacterial- and plant-type PEPC can interact with each other to form a hetero-octameric complex that is less sensitive to inhibition by malate^32^. Although the functional and physiological implications await future studies, the unusual involvement of bacterial-type PEPC speaks to the uniqueness of *Isoetes*’ underwater CAM.

### No evidence for convergent evolution of PEPC

Plant-type PEPC was recently shown to undergo convergent amino acid substitutions in concert with the evolution of CAM^10^. An aspartic acid (D) residue appears to have been repeatedly selected across multiple origins of CAM such as in *K. fedtschenkoi* and *P. equestris*, although notably not in *A. comosus*^10^. This residue is situated near the active site, and based on *in vitro* assays, the substitution to aspartic acid significantly increased PEPC activity^10^. However, in *I. taiwanensis* we did not observe the same substitution in any copies of PEPC (Fig. 4); instead, they have arginine (R) or lysine (H) at this position like PEPC from many non-CAM plants. This lack of sequence convergence between *Isoetes* and flowering plants could be the result of their substantial phylogenetic distance and highly divergent life histories. Alternatively, it is also likely that the substitution is relevant only in the context of plant-type PEPC, and as *I. taiwanensis* recruited the bacterial-type PEPC, the aspartic acid residue might not serve the same purpose.

**Fig. 4:**
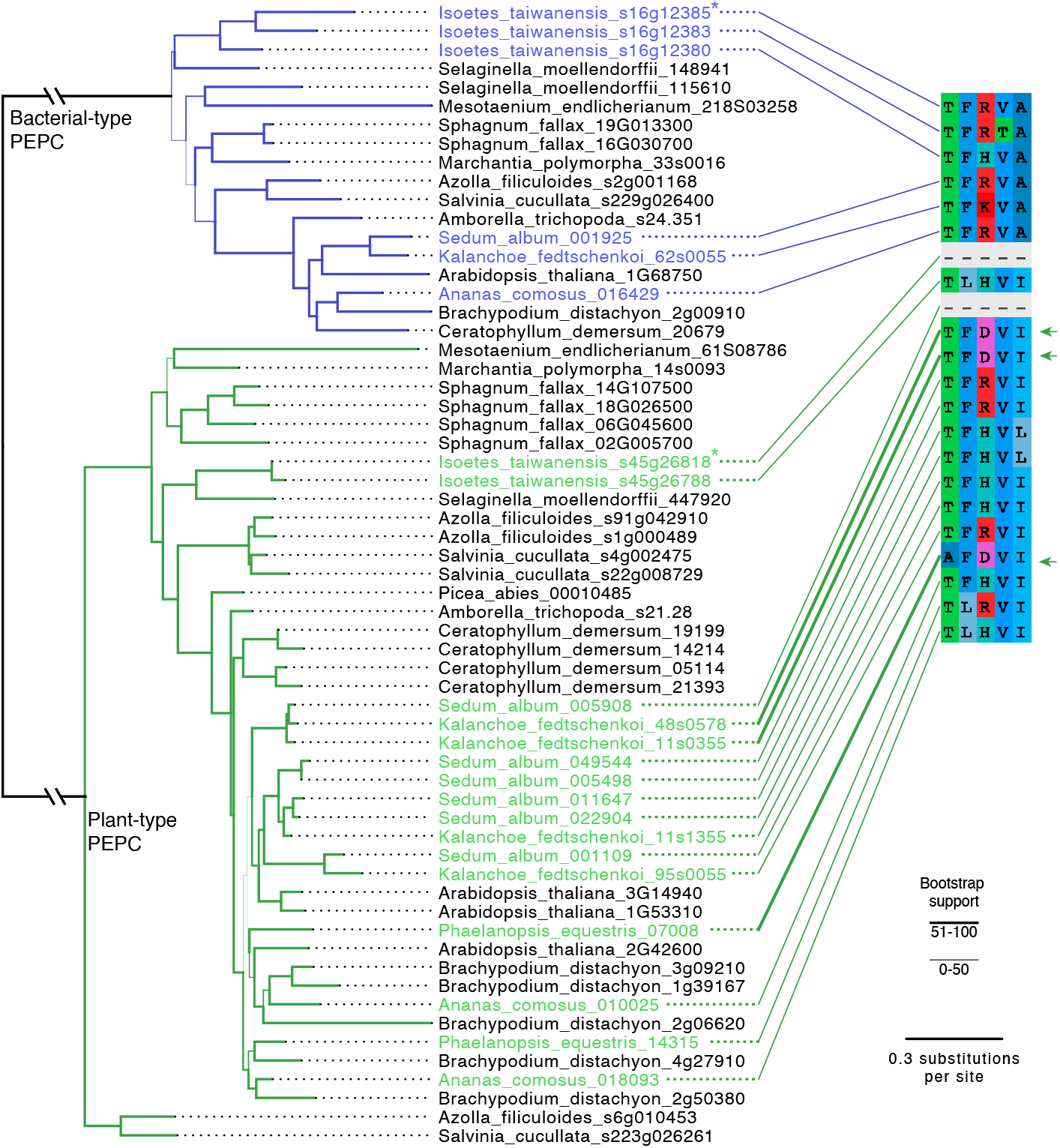
A lack of PEPC sequence convergence in *taiwanensis*. Copies with putative convergent amino acid sequence (D at position 3 in alignment) are indicated by thickened connecting lines and green arrows. Copies of bacterial-type and plant-type PEPC shown to cycle in *I. taiwanensis* are marked with asterisks (*). Branch thickness indicates bootstrap support.

### A unique circadian regulation in Isoetes

Previous analysis of the *A. comosus* genome found promoter regions of multiple key CAM pathway genes containing known circadian cis-regulatory elements (CREs) including Morning Element (ME: CCACAC), Evening Element (EE: AAATATCT), CCA1-binding site (CBS: AAAAATCT), G-box (CACGTG) and TCP15-binding motif (NGGNCCCAC)^9^. This suggests that expression of CAM genes in pineapple is largely under the control of a handful of known circadian clock elements. The direct involvement of circadian CREs was corroborated by a later study of the facultative CAM plant *S. album* where shifts in diel expression patterns were tied to a shift in TOD-specific enrichment of CREs: EE and Telobox (TBX: AAACCCT)^8^.

In order to examine the role of the circadian clock and light/dark cycles in regulating *I. taiwanensis* CAM, we used the HAYSTACK pipeline^33^ to identify all genes with TOD expression patterns. We predicted 3,241 cycling genes, which is 10% of the expressed genes. While 10% is low compared to land plants that have been tested under this condition (LDHH)— usually at 30-50% genes^8,33,34^, a recent study found a reduced number of cycling genes in another aquatic plant *Wolffia australiana* (duckweed/watermeal)^35^. Accordingly, decreased cycling may be a feature of aquatic plants.

Core circadian clock genes such as *LATE ELONGATED HYPOCOTYL* (*LHY*; Fig. 5a), *PSEUDO-RESPONSE REGULATOR 7* (*PRR7*), *LUX ARRHYTHMO* (*LUX*), and *EARLY FLOWERING 3* (*ELF3*) (Supplementary Figure S27), cycle with the expected TOD expression seen in their *Arabidopsis* orthologs^33^. However, *ZEITLUPE* (*ZTL*) does not appear to cycle in *I. taiwanensis*, in contrast to orthologues in *Arabidopsis* and *Selaginella*^36^. Furthermore, *TIMING OF CAB2 1/PSEUDO-RESPONSE REGULATOR 1* (*TOC1/PRR1*) and *GIGANTEA (GI)*, which are typically single-copy genes in land plants, have respectively 3 and 5 predicted genes in distinct genomic locations; similarly an increased number of homologs was found in the facultative CAM plant *S. album*^8^. Closer inspection confirmed all 3 *TOC1/PRR1* paralogs are full length, while only 1 of the *GI* genes (*GIa*) is full length and 1 other (*GIb*) is a true partial/truncated (and expressed) paralog. Surprisingly, all 3 copies of *TOC1/PRR1* have dawn-specific expression compared to the dusk-specific expression found in all plants tested to date^37^ (Fig. 5b).

**Fig. 5:**
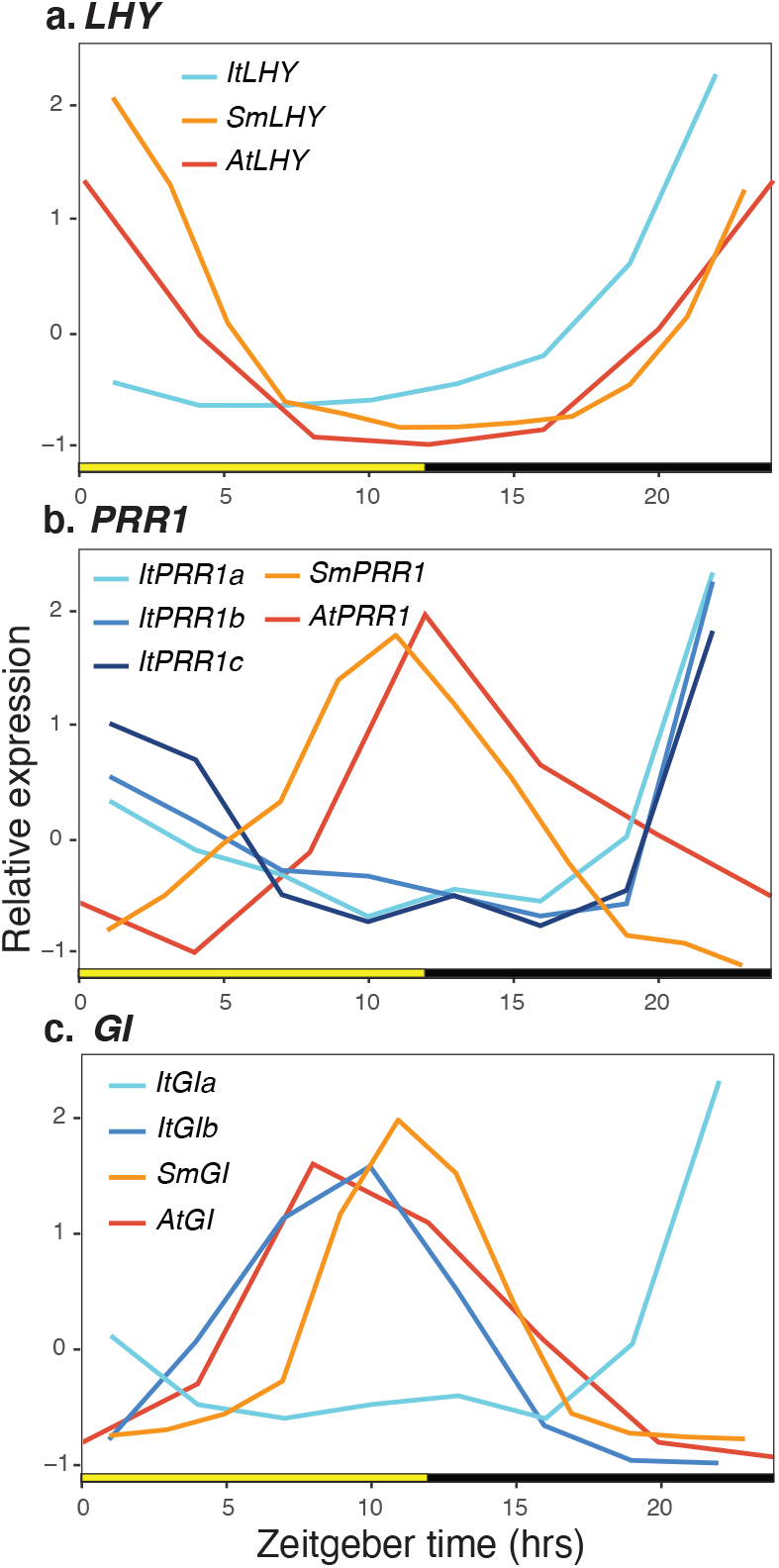
Expression of key circadian associated genes is shifted in *I. taiwanensis*. **a**, *LATE ELONGATED HYPOCOTYL (LHY)* (*CCA1*), **b**, *PSEUDO-RESPONSE REGULATOR 1* (*PRR1*), and **c**, *GIGANTEA* (*GI*) orthologs in *Isoetes* (blue lines), *Selaginella* (orange line) and *Arabidopsis* (red line) over the day. Day (yellow box); Night (black box); Zeitgeber time (ZT) is the number of hours (hrs) after lights-on (0 hrs).

In addition, *GIa* and *GIb* have antiphasic expression, with the full length *GIa* having dusk-specific expression, which is consistent with other plants, and *GIb* having dawn-specific expression (Fig. 5c).

The duplications and divergent expression patterns of *TOC1/PRR1* and *GI* in *I. taiwanensis* have important implications on circadian clock evolution. Despite the TOD expression of core circadian clock genes being highly conserved since the common ancestor of green algae and angiosperms, the mechanisms may be simpler in algae^38^ and mosses^39^. This is largely due to a lack of key components of the evening-phased loop including *PRR1, GI*, and *ZTL* in *P. patens* and the absence of the same along with morning-phased loop genes *ELF3* and *ELF4* in algae^36^. While *I. taiwanensis* possesses all the major clock genes that are found in other vascular plants, lineage specific expansion and phase-shifted gene expression in the evening-phased loop could indicate that circadian control was less conserved during the early evolution of land plants. However, *Selaginella* exhibits very similar expression of various circadian modules relative to other vascular plants and likewise, possesses a single copy of both *GI* and *PRR1*^36^. It is thus possible that the unique TOD architecture in *I. taiwanensis* represents a more recent adaptation to its aquatic CAM lifestyle. As a comparison, *S. album* similarly has multiple duplicated clock genes and its transition to CAM is associated with significant shifts in both phase and amplitude of gene expression^8^. To further investigate the relationship between clock and CAM in *I. taiwanensis*, we next focused on characterizing the circadian CREs.

### Canonical circadian CREs are not enriched in Isoetes CAM cycling genes

We used ELEMENT^33^ to exhaustively search the promoter region of cycling genes for putative CRE motifs. Following *de novo* identification, putative CREs were compared to known transcription factor binding sites in *Arabidopsis* to determine to what degree their functions might be conserved between *Isoetes* and flowering plants. We identified 16 significantly enriched CREs motifs in the 500 bp 5’ promoter region of cycling genes identified by HAYSTACK, and clustered them according to TOD expression (Supplementary Table S3). Half of the motifs shared some degree of sequence similarity to known circadian CREs previously identified in *Arabidopsis*, including the EE as well as two ‘ACGT’-containing elements (Gbox-like) and two TBX-containing motifs^33^. In the case of TBX, both motifs were associated with peak expression at dusk (at around 12 hrs after lights on; Zeitgeber Time [ZT]) in *I. taiwanensis* (Fig. 6a,b), similar to *Arabidopsis* under light/dark cycles alone^33^. On the other hand, the EE appear to be associated with peak expression at different TOD. In *Arabidopsis*, the EE is enriched in genes with peak expression at dusk (ZT = 12), but in *I. taiwanensis*, this pattern is shifted, with the EE associating with genes that peak in expression around mid-day (ZT= 6) (Fig. 6c). Additionally, while the two ‘ACGT’-containing elements were found upstream of genes that exhibited significant cycling behavior, neither was strongly associated with peak expression at a particular TOD. We also found an unidentified CRE (AGAATAAG) strongly associated with peak expression in the morning (ZT = 4)(Fig. 6d).

**Fig. 6:**
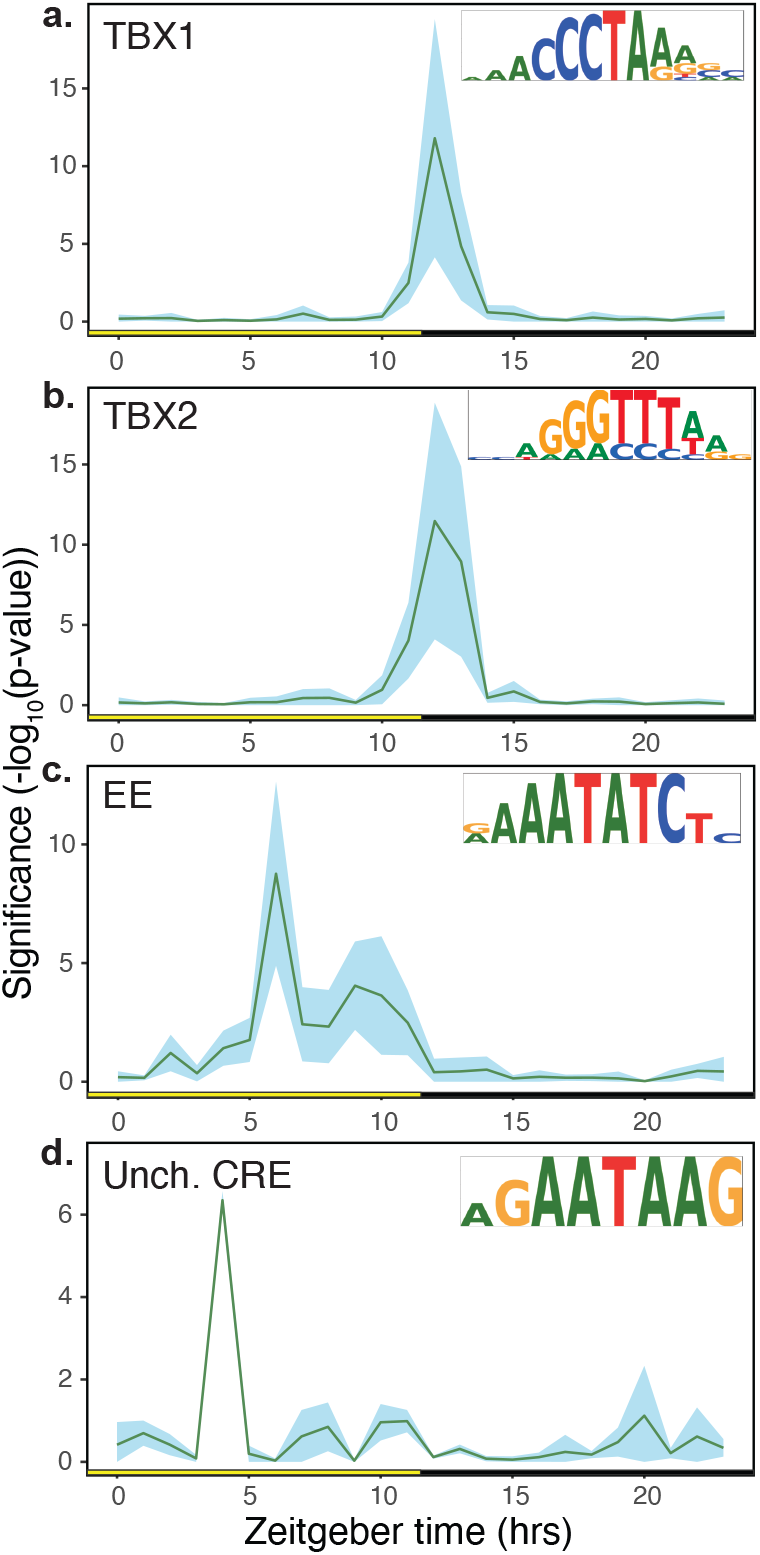
Multiple CREs exhibit time-structured enrichment in *I. taiwanensis*. **a**,**b**, Two telobox (TBX) containing motifs showed similar patterns to one another, both being enriched in genes with peak expression at dusk. **c**, A motif containing Evening Element (EE) was significantly enriched in genes with peak expression at mid-day. **d**, A novel motif was significantly enriched at mid-day as well. Day (yellow box); Night (black box); Zeitgeber time (ZT) is the number of hours (hrs) after lights on (0 hrs).

We next examined the connection between circadian CREs and CAM genes in *I. taiwanensis*. Interestingly, with the exception of the RVE1/2 motif, we did not find significant enrichment of any known circadian CREs in CAM cycling genes relative to non-cycling paralogues. While a targeted search of CAM cycling gene promoters did uncover circadian CREs including the CBS, TCP15, TBX, and EE (Supplementary Table S4), none were strongly associated with either light or dark phase CAM gene expression. In addition, both ME and G-box were conspicuously absent from the promoter regions of cycling CAM photosynthetic genes.

In sum, TOD-specific enrichment of CREs appears to differ significantly from *Arabidopsis*. While some CRE sequences themselves are conserved between lycophytes and angiosperms, their interaction with various transcription factors and subsequent regulatory function could be quite different in *Isoetes*. Importantly, our results stand in contrast to other CAM plants such as *S. album*^8^ and *A. comosus*^9^ where CAM genes appeared to be under the direct control of a handful of strictly conserved circadian CREs. These results either suggest that the circadian clock network that emerged in *Isoetes*, which included the addition of central components *GI* and *PRR1*, was quite different than that found to be highly conserved in seed plants, or there is significant TOD innovation associated with the evolution of underwater CAM. Additional *Isoetes* genomes and TOD analysis of underwater CAM plants will be required to narrow these hypotheses.

### Conclusion

The assembly and analyses of the *I. taiwanensis* genome bridges a substantial gap in our knowledge of vascular plant evolution. We have combined genomic and transcriptomic data to corroborate one of the two hypothesized WGDs in *Isoetes* relative to its closest extant relative *Selaginella*, highlighting the contrasting history of WGD in these two lineages. Importantly, comparison of TOD gene expression with genomic sequence data has given us unique insights into the convergent evolution of CAM photosynthesis, not only in a lycophyte, but also in the aquatic environment. As such, our analysis stands as a necessary counterpoint to similar studies previously conducted in terrestrial angiosperms. Shifts in expression of CAM pathway genes and the recruitment of bacterial-type PEPC in *I. taiwanensis* demonstrate a remarkable degree of plasticity in the convergent evolution of this complex trait throughout vascular plants. Likewise, differences in the enrichment of CREs associated with circadian gene expression suggest that control of CAM, as well as other processes tied to the circadian clock, may have diverged significantly since the common ancestor of *Isoetes* and flowering plants. We propose that the emergence of underwater CAM may have followed a distinct route in *Isoetes*, shedding new light on a classic example of convergent evolution of a complex plant trait.

## Methods

### Plant sample

*Isoetes taiwanensis* is endemic to a small pond in Northern Taiwan and has been *ex situ* propagated in Taiwan Forestry Research Institute. This species is expected to have a low genetic diversity due to a very restricted distribution and a small population size. The voucher specimen was deposited at TAIF herbarium.

### Genome size estimate

The genome size of *I. taiwanensis* was first determined by flow cytometry following the protocols outlined in Kuo et al.^40^ and Li et al^28^. The flow cytometric experiments were performed on BD FACSCan system (BD Biosciences, USA), and the Beckman buffer^41^ was used with 0.5% (v/v) 2-mercaptoethanol, 40 mg ml^-1^ PVP-40, and 0.1 mg ml^-1^ RNaseA added. We used *Zea mays* (1C = 5.57pg^42^) as the internal standard. To confirm the flow cytometry-based measurement, a K-mer frequency distribution was generated from Illumina 2×150 bp paired reads (described below) using Jellyfish^43^, which was then input into GenomeScope^44^ and an inhouse pipeline to estimate genome size and heterozygosity.

### Genome sequencing

High molecular weight (HMW) DNA was extracted using a modified CTAB method on isolated nuclei. First, leaf tissues were ground in liquid nitrogen, and the powder was resuspended in the Beckman buffer (same as in our flow cytometric experiments). We then used 30µm nylon circular filters (Partec, Germany) to remove tissue debris, and precipitated nuclei with 100g centrifugation under 4°C for 20 minutes. For the downstream CTAB procedures, we followed the protocol outlined in Kuo^45^. HMW DNA was QC’d on an agarose gel for length and quantified on a bioanalyzer. Unsheared HMW DNA was used to make Oxford Nanopore Technologies (ONT) ligation-based libraries (Oxford, UK). Libraries were prepared starting with 1.5ug of DNA and following all other steps in ONT’s SQK-LSK109 protocol. Final libraries were loaded on an ONT flowcell (v9.4.1) and run on the GridION. Bases were called in real-time on the GridION using the flip-flop version of Guppy (v3.1). The resulting fastq files were concatenated and used for downstream genome assembly steps. The same batch of HMW genomic DNA was used to construct Illumina (Illumina, USA) libraries for estimating genome size (above) and correcting residual errors in the ONT assembly. Libraries were constructed using the KAPA HyperPrep Kit (Kapa Biosystems, Switzerland) followed by sequencing on an Illumina NovaSeq6000 with 2×150 bp paired-ends.

### Genome assembly

ONT reads were assembled using minimap2 and miniasm^46^, and the resulting draft assembly was then polished by racon^47^ (with nanopore reads) and pilon^48^ (with Illumina reads). Because the plants were grown non-axenically under water, the assembly inevitably contained contaminations. We therefore used blobtools^49^ to identify non-plant contigs based on a combination of contig read coverage, taxonomic assignment, and GC content.

To further scaffold the assembly, we generated a genome map using Bionano with the Direct Label and Stain chemistry and DLE-1 labeling. For this, high molecular weight DNA was extracted using the Bionano Plant DNA Isolation Kit. Hybrid scaffolding, combining the nanopore draft and Bionano map, was done on the Bionano Saphyr computing platform at the McDonnell Genome Institute at Washington University. We then gap-filled the scaffolded genome using two rounds of LR_Gapcloser^50^ (3 iterations each and a pilon polishing in between. Finally, to remove redundancy the purge_haplotigs pipeline^51^ was used to obtain the v1 assembly. The circular chloroplast genome was assembled from Illumina data using the GetOrganelle^52^ toolkit.

### Repeat annotation

We generated a custom *I. taiwanensis*-specific repeat library using LTR-retriever^53^ and RepeatModeler^54^. To identify and remove repeats with homology to plant proteins, we used BLASTx to query each repeat against the uniprot plant protein database (e-value threshold at 1e-10). The resulting library was then input into RepeatMasker^55^ to annotate and mask the repetitive elements in the *I. taiwanensis* genome.

### Gene annotation

We trained two *ab initio* gene predictors, AUGUSTUS^56^ and SNAP^57^, on the repeat-masked genome using a combination of protein and transcript evidence. For the protein evidence, we relied on the annotated proteomes from *Selaginella moellendorffii*^16^ and *S. lepidophylla*^18^, and for the transcript evidence, we used the RNA-seq data from our time-course experiment and a separate corm sample. To train AUGUSTUS, BRAKER2^58^ was used and the transcript evidence was input as an aligned bam file. On the other hand, SNAP was trained under MAKER with 3 Iterations, and in this case, the transcript evidence was supplied as a *de novo* assembled transcriptome done by Trinity^59^. After AUGUSTUS and SNAP were trained, they were fed into MAKER^60^ along with all the evidence to provide a synthesized gene prediction. Gene functional annotation was done using the eggNOG-mapper v2^61^. To filter out spurious gene models, we removed genes that met none of the following criteria: (1) a transcript abundance greater than zero in any sample (as estimated by Stringtie^62^), (2) has functional annotation from eggNOG, and (3) was assigned into orthogroups in an Orthofinder^20^ run (see below). The resulting gene set was used in all subsequent analyses.

### Homology assessment and gene family analysis

Homology was initially assessed with Orthofinder^20^ using genomic data from a range of taxa from across the plant tree of life including all CAM plant genomes published to date: *Amborella trichopoda*^63^, *Ananas comosus*^9^, *Anthoceros agrestis*^24^, *Arabidopsis thaliana*^64^, *Azolla filiculoides*^28^, *Brachypodium distachyon*^65^, *Ceratophyllum demersum*^66^, *Isoetes taiwanensis* (this study), *Kalanchoe fedtschenkoi*^10^, *Marchantia polymorpha*^23^, *Medicago truncatula*^67^, *Nelumbo nucifera*^68^, *Nymphaea colorata*^69^, *Phalaenopsis equestris*^11^, *Physcomitrium patens*^22^, *Picea abies*^70^, *Salvinia cucullata*^28^, *Sedum album*^8^, *Selaginella moellendorffii*^16^, *Sphagnum fallax* (*Sphagnum fallax* v0.5, DOE-JGI, http://phytozome.jgi.doe.gov/), *Spirodela polyrhiza*^71^, *Utricularia gibba*^72^, *Vitus vinifera*^73^, and *Zostera marina*^74^, and one algal genome: *Mesotaenium endlicherianum*^75^. Following homology assessment, the degree of overlap between gene families was assessed using the UpsetR^76^ package in R.

### RNA editing analysis

RNA-seq data was first mapped to combined nuclear and chloroplast genome assemblies using HISAT2^77^. The reads mapping to the chloroplast genome were extracted using samtools^78^. SNPs were called using the mpileup function in bcftools^79^. The resulting vcf files were filtered using bcftools to remove samples with a depth < 20, quality score < 20 and mapping quality bias < 0.05. After filtering, C-to-U and U-to-C edits were identified using an alternate allele frequency threshold of 10%. Finally, RNA editing sites were related to specific genes using the intersect command in bedtools^80^ and characterized using a custom python script.

### Ks analysis

Ks divergence was calculated by several different methods. Initially, a whole paranome Ks distribution was generated using the ‘wgd mcl’ tool^81^. Self-synteny was then assessed in i-Adhore and Ks values were calculated and plotted for syntenic pairs only using the ‘wgd syn’ tool^81^. To conduct Ks analysis of related species, RNA-seq data was downloaded from the SRA database for *Isoetes yunguiensis, I. sinensis, I. drummondii, I. echinospora, I. lacustris* and *I. tegetiformans*. Transcriptomes were assembled using SOAPdenovo-Trans^82^ with a Kmer length of 31. Next, for each *Isoetes* genome and transcriptome, we used the DupPipe pipeline to construct gene families and estimate the age distribution of gene duplications^83,84^. We translated DNA sequences and identified ORFs by comparing the Genewise^85^ alignment to the best-hit protein from a collection of proteins from 25 plant genomes from Phytozome^86^. For all DupPipe runs, we used protein-guided DNA alignments to align our nucleic acid sequences while maintaining the ORFs. We estimated Ks divergence using PAML^87^ with the F3×4model for each node in the gene family phylogenies. We then used mixture modeling to identify significant peaks consistent with a potential WGD and to estimate their median paralog Ks values. Significant peaks were identified using a likelihood ratio test in the boot.comp function of the package mixtools^88^ in R.

### Estimation of orthologous divergence

To place putative WGDs in relation to lineage divergence, we estimated the synonymous divergence of orthologs among pairs of species that may share a WGD based on their phylogenetic position and evidence from the within-species Ks plots. We used the RBH Orthologue pipeline^84^ to estimate the mean and median synonymous divergence of orthologs, and compared those with the synonymous divergence of inferred paleopolyploid peaks. We identified orthologs as reciprocal best blast hits in pairs of transcriptomes. Using protein-guided DNA alignments, we estimated the pairwise synonymous divergence for each pair of orthologs using PAML^87^ with the F3X4 model.

### Phylogenetic assessment of ancient whole genome duplication

WGD inference was conducted by phylogenomic reconciliation using the WhALE package implemented in Julia^27^. First, prior to WhALE analysis, Orthofinder^20^ was used to identify groups of orthologous genes among 7 species representing 3 taxonomic groups (bryophytes, lycophytes, and ferns): *Azolla filiculoides*^28^, *Isoetes taiwanensis* (this study), *Marchantia polymorpha*^23^, *Physcomitrium patens*^22^, *Salvinia cucullata*^28^, *Selaginella moellendorffii*^16^, and *Sphagnum fallax* (*Sphagnum fallax v0*.*5*, DOE-JGI, http://phytozome.jgi.doe.gov/). These species were chosen based on phylogenetic relatedness, availability of a high-quality genome assembly, and previous assessment for the presence or absence of WGD. The resulting orthogroups were filtered using a custom python script to remove the 5% largest orthogroups and those with less than 3 taxa. Additionally, WhALE requires removal of gene families that do not contain at least one gene in both bryophytes and ferns to prevent the inclusion of gene families originating after divergence from the most recent common ancestor. Alignments were generated for the filtered orthogroups in PRANK^89^ using the default settings. A posterior distribution of trees was obtained for each gene family in MrBayes 3.2.6^90^ using the LG model. Chains were sampled every 10 generations for 100,000 generations with a relative burn-in of 25%. Following the Bayesian analysis, conditional clade distributions (CCDs) were determined from posterior distribution samples using ALEobserve in the ALE software suite^91^. CCD files were subsequently filtered using the ccddata.py and ccdfilter.py scripts provided with the WhALE program. A dated, ultrametric species tree was generated using the ‘ape’ package in R^92^, in which branch lengths were constrained according to 95% highest posterior density of ages, assuming that bryophytes are monophyletic, as reported by Morris et al.^93^. Finally, the filtered CCD files were loaded in Julia along with the associated species phylogeny. A hypothetical WGD node was inferred at 200 million years ago (MYA) along the branch leading to *I. taiwanensis*, prior to the estimated crown age of extant *Isoetes*^94^. Modifying the hypothetical age of this WGD node did not affect the outcome. Additional WGD nodes were placed as positive controls along branches leading to *Physcomitrium patens* and *Azolla filiculoides* at 40 MYA and 60 MYA, respectively, based on previous studies^22,28^. A false WGD event was also placed arbitrarily in *Marchantia polymorpha* at 160 MYA as a negative control. A WhALE ‘problem’ was constructed using an independent rate prior and MCMC analysis was conducted using the DynamicHMC library in Julia (https://github.com/tpapp/DynamicHMC.jl) with a sample size of 1000.

### Phylogenetic analysis of root, stomata, and CAM pathway genes

Following clustering of homologs in Orthofinder, we conducted phylogenetic analysis of several gene families of interest, including those containing *SMF, FAMA, TMM, RSL*, and *PEPC* genes, were subsequently identified based on homology using gene annotations from *Arabidopsis*. Gene trees from Orthofinder were initially used to identify paralogues and remove fragmented genes where appropriate. In the case of *PEPC*, orthogroups containing “bacterial-type” and “plant-type” *PEPC* were combined prior to alignment. Next, amino acid sequences were aligned using MUSCLE^95^ under default settings and trimmed using TrimAL with the -strict flag. An amino acid substitution model was selected according to the Bayesian Information Criterion (BIC) in ModelFinder^96^ prior to phylogenetic reconstruction by maximum likelihood in IQ-TREE v1.6.12^97^ with 1000 ultrafast^98^ bootstrap replicates.

### Phylogenetic analysis of genes salient to the phenylpropanoid and lignin biosynthesis pathway

The datasets used for phylogenetic analysis were based on de Vries et al.^99^ with added *I. taiwanensis* sequences. In brief, we assembled a dataset of predicted proteins from (A) the genomes of seventeen land plants: *Anthoceros agrestis* as well as *Anthoceros punctatus*^24^, *Amborella trichopoda*^63^, *Arabidopsis thaliana*^64^, *Azolla filiculoides*^28^, *Brachypodium distachyon*^65^, *Capsella grandiflora*^100^, *Gnetum montanum*^101^, *Isoetes taiwanensis* (this study), *Marchantia polymorpha*^23^, *Nicotiana tabacum*^102^, *Oryza sativa*^103^, *Physcomitrium patens*^22^, *Picea abies*^70^, *Salvinia cucullata*^28^, *Selaginella moellendorffii*^16^, and *Theobroma cacao*^104^; (B) the genomes of seven streptophyte algae: *Chlorokybus atmophyticus*^105^, *Chara braunii*^106^, *Klebsormidium nitens*^107^, *Mesotaenium endlicherianum*^75^, *Mesostigma viride*^105^, *Penium margaritaceum*^108^, *Spirogloea muscicola*^75^—additionally, we included sequences found in the transcriptomes of *Spirogyra pratensis*^109^, *Coleochaete scutata* as well as *Zygnema circumcarinatum*^110^, and *Coleochaete orbicularis*^111^; (C) the genomes of eight chlorophytes: *Bathycoccus prasinos*^112^, *Chlamydomonas reinhardtii*^113^, *Coccomyxa subellipsoidea*^114^, *Micromonas* sp. as well as *Micromonas pusilla*^115^, *Ostreococcus lucimarinus*^116^, *Ulva mutabilis*^117^, *Volvox carteri*^118^. For phenylalanine ammonia-lyase, additional informative sequences were added based on de Vries et al.^119^.

Building on the alignments published in de Vries et al.^99^, homologs of each gene family (detected in the aforementioned species via BLASTp) were (re-)aligned using MAFFT v7.475^120^ with a L-INS-I approach; both full and partial sequences from *I. taiwanensis* were retained. We constructed maximum likelihood phylogenies using IQ-TREE 2.0.6^121^; 1000 ultrafast^98^ bootstrap replicates were computed. To determine the best model for protein evolution, we used ModelFinder^96^ and picked the best models based on BIC. Residue information was mapped next to the tree based on structural analyses by Hu et al.^122^, Pan et al.^123^, Louie et al.^124^, Youn et al.^125^ and Ferrer et al.^126^.

### Time course titratable acidity and RNA-seq experiments

Leaves of *I. taiwanensis* were taken from five individuals every 3 hours over a 27-hour period on a 12-hour light/dark cycle and constant temperature. To measure changes in acidity over time, a portion of the leaf tissues was weighed, mixed with 3.5-5ml of ddH20, and titrated with 0.0125M NaOH solution until pH = 7.0. At the same time, we froze the leaf tissues in liquid nitrogen, and extracted RNA using a modified CTAB protocol^127^. RNA quality was examined on a 1% agarose gel and RNA concentration was quantified using the Qubit RNA HS assay kit (Invitrogen, USA). 2ug of total RNA was used to construct stranded RNA-seq libraries using the Illumina TruSeq stranded total RNA LT sample prep kit (RS-122-2401 and RS-122-2402). Multiplexed libraries were pooled and sequenced on an Illumina NovaSeq6000 with 2×150 bp paired-ends.

### Differential expression analysis

RNA-seq reads were mapped to the combined nuclear and chloroplast genome using HISAT2^77^. Stringtie^62^ was used to assemble transcripts and estimate transcript abundance. A gene count matrix was produced using the included prepDE.py script. We imported gene count data into the DESEQ2 package in R^128^ for read normalization using its median of ratios method as well as identification and removal of outlier samples using multidimensional scaling. A single outlier sample from each of six time points (1hr, 4hrs, 7hrs, 10hrs, 13hrs and 19hrs) was removed from the final dataset. The resulting dataset was used to analyze temporal gene expression patterns in the R package maSigPro^129^. Using maSigPro, genes with significantly differential expression profiles were identified by computing a regression fit for each gene and filtered based on the associated p-value (p<0.001).

### HAYSTACK global cycling prediction

Genes with mean expression across all the time points below 1 TPM were considered “not expressed” and filtered prior to cycling prediction with HAYSTACK (https://gitlab.com/NolanHartwick/super_cycling)^33^. HAYSTACK operates by correlating the observed expression levels of each gene with a variety of user specified models that represent archetypal cycling behavior. We used a model file containing sinusoid, spiking traces, and various rough linear interpolations of sinusoids with periods ranging from 20 hours to 28 hours in one-hour increments and phases ranging from 0-23 hours in one-hour increments. Genes that correlated with their best fit model at a threshold of R > 0.8 were classified as cyclers with phase and period defined by the best fit model. This threshold for calling cycling genes is based on previous validated observations^8,33,34,130^. We also validated this threshold by looking at the cycling of known circadian clock genes (Fig. 5).

### ELEMENT cis-regulatory elements analysis

Once cycling genes in I. *taiwanensis* were identified, we were able to find putative cis-acting elements associated with TOD expression. Promoters, defined as 500 bp upstream of genes, were extracted for each gene and processed by ELEMENT (https://gitlab.com/salk-tm/snake_pip_element)^33,131,132^. Briefly, ELEMENT generates an exhaustive background model of all 3-7 K-mer using all of the promoters in the genome, and then compares the K-mers (3-7 bp) from the promoters for a specified gene list. Promoters for cycling genes were split according to their TOD expression into “phase” gene lists and K-mers that were overrepresented in any of these 24 promoter sets were identified by ELEMENT. By splitting up cycling genes according to their associated phase, we gained the power to identify K-mers associated with TOD-specific cycling behavior at every hour over the day. Our threshold for identifying a K-mer as being associated with cycling was an FDR < 0.05 in at least one of the comparisons. The significant K-mers were clustered according to sequence similarity (Fig. 6).

### Promoter motif identification

Core CAM genes with significantly differential diel expression profiles (as identified in maSigPro) including *β-CA, PEPC, PEPCK, ME, MDH*, and *PPDK* were selected for motif enrichment analysis. Enriched motifs were identified relative to a background consisting of non-cycling paralogues of photosynthetic genes using the AME utility^133^. Promoters were searched for known motifs from the *Arabidopsis* promoter binding motif database^134^ with FIMO^135^.

## Data availability

All the raw sequences were deposited in the NCBI Sequence Read Archive under the BioProject PRJNA735564. Genome assembly and annotation are available at https://genomevolution.org/coge/GenomeInfo.pl?gid=61511. Sequence alignments and tree files can be found at https://github.com/dawickell/Isoetes_CAM.

## Acknowledgements

The authors would like to thank Karolina Heyduk, Peter Schafran, and Arthur Zwaenepoel for their advice and support to various aspects of this project. J.d.V. is supported through funding from the European Research Council (ERC) under the European Union’s Horizon 2020 research and innovation programme (grant no. 852725; ERC Starting Grant ‘TerreStriAL’). A.D.A. and A.D. are grateful for being supported through the International Max Planck Research School (IMPRS) for Genome Science.

## Author contributions

D.W., L.-Y.K., T.P.M. and F.-W.L. coordinated the project. Y.-M.H. provided the plant materials. L.-Y.K. carried out the time-course experiment and nucleic acid extraction. T.P.M. and F.-W.L. sequenced and assembled the genome. H.-P.Y. and F.-W.L. annotated the genome. D.W. assembled the plastome and profiled RNA-editing. D.W. circumscribed gene families and examined genes related to stomata and root development. A.D.A., I.I., A.M., S.d.V. and J.d.V. characterized lignin biosynthesis genes. D.W., Z.L. and M.S.B. carried out WGD analysis. D.W. analyzed expressions of CAM pathway genes. N.T.H. and T.P.M. carried out HAYSTACK and ELEMENT analyses. D.W., T.P.M. and F.-W.L. synthesized and wrote the manuscript.

## References

1. Pteridophyte Phylogeny Group I. A community-derived classification for extant lycophytes and ferns. J. Syst. Evol. 54, 563–603 (2016).

2. Pigg, K. B. Isoetalean lycopsid evolution: from the Devonian to the present. Am. Fern J. 91, 99–114 (2001).

3. Keeley, J. E. Distribution of diurnal acid metabolism in the genus Isoetes. Am. J. Bot. 69, 254–257 (1982).

4. Keeley, J. E. CAM photosynthesis in submerged aquatic plants. Bot. Rev. 64, 121–175 (1998).

5. Aulio, K. Differential expression of diel acid metabolism in two life forms of Littorella uniflora (l.) Aschers. New Phytologist 100, 533–536 (1985).

6. Suissa, J. S. & Green, W. A. CO2 starvation experiments provide support for the carbon-limited hypothesis on the evolution of CAM-like behaviour in Isoëtes. Ann. Bot. 127, 135– 141 (2021).

7. Keeley, J. E. Isoetes howellii: A submerged aquatic cam plant? Am. J. Bot. 68, 420–424 (1981).

8. Wai, C. M. et al. Time of day and network reprogramming during drought induced CAM photosynthesis in Sedum album. PLOS Genetics 15, e1008209 (2019).

9. Ming, R. et al. The pineapple genome and the evolution of CAM photosynthesis. Nat. Genet. 47, 1435–1442 (2015).

10. Yang, X. et al. The Kalanchoë genome provides insights into convergent evolution and building blocks of crassulacean acid metabolism. Nat. Commun. 8, 1899 (2017).

11. Cai, J. et al. The genome sequence of the orchid Phalaenopsis equestris. Nat. Genet. 47, 65–72 (2015).

12. Zhang, L. et al. Origin and mechanism of crassulacean acid metabolism in orchids as implied by comparative transcriptomics and genomics of the carbon fixation pathway. Plant J. 86, 175–185 (2016).

13. Heyduk, K. et al. Altered Gene Regulatory Networks Are Associated With the Transition From C3 to Crassulacean Acid Metabolism in Erycina (Oncidiinae: Orchidaceae). Front. Plant Sci. 9, 2000 (2018).

14. Heyduk, K. et al. Shared expression of crassulacean acid metabolism (CAM) genes pre-dates the origin of CAM in the genus Yucca. J. Exp. Bot. 70, 6597–6609 (2019).

15. Abraham, P. E. et al. Transcript, protein and metabolite temporal dynamics in the CAM plant Agave. Nat. Plants 2, 16178 (2016).

16. Banks, J. A. et al. The Selaginella genome identifies genetic changes associated with the evolution of vascular plants. Science 332, 960–963 (2011).

17. Xu, Z. et al. Genome analysis of the ancient tracheophyte Selaginella tamariscina reveals evolutionary features relevant to the acquisition of desiccation tolerance. Mol. Plant 11, 983–994 (2018).

18. VanBuren, R. et al. Extreme haplotype variation in the desiccation-tolerant clubmoss Selaginella lepidophylla. Nat. Commun. 9, 13 (2018).

19. One Thousand Plant Transcriptomes Initiative. One thousand plant transcriptomes and the phylogenomics of green plants. Nature 574, 679–685 (2019).

20. Emms, D. M. & Kelly, S. OrthoFinder: phylogenetic orthology inference for comparative genomics. Genome Biol. 20, 238 (2019).

21. Schmutz, J. et al. Genome sequence of the palaeopolyploid soybean. Nature 463, 178–183 (2010).

22. Lang, D. et al. The Physcomitrella patens chromosome-scale assembly reveals moss genome structure and evolution. Plant J. 93, 515–533 (2018).

23. Diop, S. I. et al. A pseudomolecule-scale genome assembly of the liverwort Marchantia polymorpha. Plant J. 101, 1378–1396 (2020).

24. Li, F.-W. et al. Anthoceros genomes illuminate the origin of land plants and the unique biology of hornworts. Nat. Plants 6, 259–272 (2020).

25. Szövényi, P., Gunadi, A. & Li, F.-W. Charting the genomic landscape of seed-free plants. Nat. Plants 7, 554–565 (2021).

26. Li, Z. & Barker, M. S. Inferring putative ancient whole-genome duplications in the 1000 Plants (1KP) initiative: access to gene family phylogenies and age distributions. Gigascience 9, (2020).

27. Zwaenepoel, A. & Van de Peer, Y. Inference of Ancient Whole-Genome Duplications and the Evolution of Gene Duplication and Loss Rates. Mol. Biol. Evol. 36, 1384–1404 (2019).

28. Li, F.-W. et al. Fern genomes elucidate land plant evolution and cyanobacterial symbioses. Nat. Plants 4, 460–472 (2018).

29. Sánchez, R. & Cejudo, F. J. Identification and expression analysis of a gene encoding a bacterial-type phosphoenolpyruvate carboxylase from Arabidopsis and rice. Plant Physiol. 132, 949–957 (2003).

30. Deng, H. et al. Evolutionary history of PEPC genes in green plants: Implications for the evolution of CAM in orchids. Mol. Phylogenet. Evol. 94, 559–564 (2016).

31. Ting, M. K. Y., She, Y.-M. & Plaxton, W. C. Transcript profiling indicates a widespread role for bacterial-type phosphoenolpyruvate carboxylase in malate-accumulating sink tissues. J. Exp. Bot. 68, 5857–5869 (2017).

32. Blonde, J. D. & Plaxton, W. C. Structural and kinetic properties of high and low molecular mass phosphoenolpyruvate carboxylase isoforms from the endosperm of developing castor oilseeds. J. Biol. Chem. 278, 11867–11873 (2003).

33. Michael, T. P. et al. Network discovery pipeline elucidates conserved time-of-day–specific cis-regulatory modules. PLoS Genet. 4, e14 (2008).

34. Filichkin, S. A. et al. Global profiling of rice and poplar transcriptomes highlights key conserved circadian-controlled pathways and cis-regulatory modules. PLoS One 6, e16907 (2011).

35. Michael, T. P. et al. Genome and time-of-day transcriptome of Wolffia australiana link morphological minimization with gene loss and less growth control. Genome Res. 31, 225–238 (2020).

36. Ferrari, C. et al. Kingdom-wide comparison reveals the evolution of diurnal gene expression in Archaeplastida. Nat. Commun. 10, 1–13 (2019).

37. Steed, G., Ramirez, D. C., Hannah, M. A. & Webb, A. A. R. Chronoculture, harnessing the circadian clock to improve crop yield and sustainability. Science 372, (2021).

38. Corellou, F. et al. Clocks in the green lineage: comparative functional analysis of the circadian architecture of the picoeukaryote Ostreococcus. Plant Cell 21, 3436–3449 (2009).

39. Holm, K., Källman, T., Gyllenstrand, N., Hedman, H. & Lagercrantz, U. Does the core circadian clock in the moss Physcomitrella patens (Bryophyta) comprise a single loop? BMC Plant Biol. 10, 1–14 (2010).

40. Kuo, L.-Y., Huang, Y.-J., Chang, J., Chiou, W.-L. & Huang, Y.-M. Evaluating the spore genome sizes of ferns and lycophytes: a flow cytometry approach. New Phytol. 213, 1974– 1983 (2017).

41. Ebihara, A. et al. Nuclear DNA, chloroplast DNA, and ploidy analysis clarified biological complexity of the Vandenboschia radicans complex (Hymenophyllaceae) in Japan and adjacent areas. Am. J. Bot. 92, 1535–1547 (2005).

42. Praça-Fontes, M. M., Carvalho, C. R., Clarindo, W. R. & Cruz, C. D. Revisiting the DNA C-values of the genome size-standards used in plant flow cytometry to choose the ‘best primary standards’. Plant Cell Rep. 30, 1183–1191 (2011).

43. Marçais, G. & Kingsford, C. A fast, lock-free approach for efficient parallel counting of occurrences of k-mers. Bioinformatics 27, 764–770 (2011).

44. Vurture, G. W. et al. GenomeScope: fast reference-free genome profiling from short reads. Bioinformatics 33, 2202–2204 (2017).

45. Kuo, L. Y. Polyploidy and biogeography in genus Deparia and phylogeography in Deparia lancea. PhD Thesis (2015).

46. Li, H. Minimap and miniasm: fast mapping and de novo assembly for noisy long sequences.Bioinformatics 32, 2103–2110 (2016).

47. Vaser, R., Sović, I., Nagarajan, N. & Šikić, M. Fast and accurate de novo genome assembly from long uncorrected reads. Genome Res. 27, 737–746 (2017).

48. Walker, B. J. et al. Pilon: an integrated tool for comprehensive microbial variant detection and genome assembly improvement. PLoS One 9, e112963 (2014).

49. Laetsch, D. R. & Blaxter, M. L. BlobTools: Interrogation of genome assemblies. F1000Res. 6, 1287 (2017).

50. Xu, G.-C. et al. LR_Gapcloser: a tiling path-based gap closer that uses long reads to complete genome assembly. Gigascience 8, (2019).

51. Roach, M. J., Schmidt, S. A. & Borneman, A. R. Purge Haplotigs: allelic contig reassignment for third-gen diploid genome assemblies. BMC Bioinformatics 19, 460 (2018).

52. Jin, J.-J. et al. GetOrganelle: a fast and versatile toolkit for accurate de novo assembly of organelle genomes. Genome Biol. 21, 1–31 (2020).

53. Ou, S. & Jiang, N. LTR_retriever: A Highly Accurate and Sensitive Program for Identification of Long Terminal Repeat Retrotransposons. Plant Physiol. 176, 1410–1422 (2018).

54. Smit, A. F. A. & Hubley, R. RepeatModeler Open-1.0. Available online at http://www.repeatmasker.org.

55. Smit, A. F. A., Hubley, R. & Green, P. RepeatMasker Open-4.0. Available online at: http://www.repeatmasker.org.

56. Stanke, M. & Morgenstern, B. AUGUSTUS: a web server for gene prediction in eukaryotes that allows user-defined constraints. Nucleic Acids Res. 33, W465–W467 (2005).

57. Korf, I. Gene finding in novel genomes. BMC Bioinformatics 5, 59 (2004).

58. Brůna, T., Hoff, K. J., Lomsadze, A., Stanke, M. & Borodovsky, M. BRAKER2: automatic eukaryotic genome annotation with GeneMark-EP+ and AUGUSTUS supported by a protein database. NAR Genom Bioinform 3, lqaa108 (2021).

59. Grabherr, M. G. et al. Full-length transcriptome assembly from RNA-Seq data without a reference genome. Nat. Biotechnol. 29, 644–652 (2011).

60. Holt, C. & Yandell, M. MAKER2: an annotation pipeline and genome-database management tool for second-generation genome projects. BMC Bioinformatics 12, 491 (2011).

61. Huerta-Cepas, J. et al. Fast Genome-Wide Functional Annotation through Orthology Assignment by eggNOG-Mapper. Mol. Biol. Evol. 34, 2115–2122 (2017).

62. Pertea, M. et al. StringTie enables improved reconstruction of a transcriptome from RNA-seq reads. Nat. Biotechnol. 33, 290–295 (2015).

63. Amborella Genome Project. The Amborella genome and the evolution of flowering plants. Science 342, 1241089 (2013).

64. Lamesch, P. et al. The Arabidopsis Information Resource (TAIR): improved gene annotation and new tools. Nucleic Acids Res. 40, D1202–10 (2012).

65. International Brachypodium Initiative. Genome sequencing and analysis of the model grass Brachypodium distachyon. Nature 463, 763–768 (2010).

66. Yang, Y. et al. Prickly waterlily and rigid hornwort genomes shed light on early angiosperm evolution. Nat Plants 6, 215–222 (2020).

67. Tang, H. et al. An improved genome release (version Mt4.0) for the model legume Medicago truncatula. BMC Genomics 15, 312 (2014).

68. Ming, R. et al. Genome of the long-living sacred lotus (Nelumbo nucifera Gaertn.). Genome Biol. 14, R41 (2013).

69. Zhang, L. et al. The water lily genome and the early evolution of flowering plants. Nature 577, 79–84 (2020).

70. Nystedt, B. et al. The Norway spruce genome sequence and conifer genome evolution. Nature 497, 579–584 (2013).

71. Wang, W. et al. The Spirodela polyrhiza genome reveals insights into its neotenous reduction fast growth and aquatic lifestyle. Nat. Commun. 5, 3311 (2014).

72. Ibarra-Laclette, E. et al. Architecture and evolution of a minute plant genome. Nature 498, 94–98 (2013).

73. Jaillon, O. et al. The grapevine genome sequence suggests ancestral hexaploidization in major angiosperm phyla. Nature 449, 463–467 (2007).

74. Olsen, J. L. et al. The genome of the seagrass Zostera marina reveals angiosperm adaptation to the sea. Nature 530, 331–335 (2016).

75. Cheng, S. et al. Genomes of Subaerial Zygnematophyceae Provide Insights into Land Plant Evolution. Cell 179, 1057–1067.e14 (2019).

76. Conway, J. R., Lex, A. & Gehlenborg, N. UpSetR: an R package for the visualization of intersecting sets and their properties. Bioinformatics 33, 2938–2940 (2017).

77. Kim, D., Paggi, J. M., Park, C., Bennett, C. & Salzberg, S. L. Graph-based genome alignment and genotyping with HISAT2 and HISAT-genotype. Nat. Biotechnol. 37, 907–915 (2019).

78. Li, H. et al. The Sequence Alignment/Map format and SAMtools. Bioinformatics 25, 2078– 2079 (2009).

79. Li, H. A statistical framework for SNP calling, mutation discovery, association mapping and population genetical parameter estimation from sequencing data. Bioinformatics 27, 2987– 2993 (2011).

80. Quinlan, A. R. & Hall, I. M. BEDTools: a flexible suite of utilities for comparing genomic features. Bioinformatics 26, 841–842 (2010).

81. Zwaenepoel, A. & Van de Peer, Y. wgd—simple command line tools for the analysis of ancient whole-genome duplications. Bioinformatics 35, 2153–2155 (2019).

82. Xie, Y. et al. SOAPdenovo-Trans: de novo transcriptome assembly with short RNA-Seq reads. Bioinformatics 30, 1660–1666 (2014).

83. Barker, M. S. et al. Multiple paleopolyploidizations during the evolution of the Compositae reveal parallel patterns of duplicate gene retention after millions of years. Mol. Biol. Evol. 25, 2445–2455 (2008).

84. Barker, M. S. et al. EvoPipes.net: Bioinformatic Tools for Ecological and Evolutionary Genomics. Evol. Bioinform. Online 6, 143–149 (2010).

85. Birney, E., Clamp, M. & Durbin, R. GeneWise and Genomewise. Genome Res. 14, 988–995 (2004).

86. Goodstein, D. M. et al. Phytozome: a comparative platform for green plant genomics. Nucleic Acids Res. 40, D1178–86 (2012).

87. Yang, Z. PAML 4: phylogenetic analysis by maximum likelihood. Mol. Biol. Evol. 24, 1586– 1591 (2007).

88. Benaglia, T., Chauveau, D., Hunter, D. & Young, D. Mixtools: An R package for analyzing finite mixture models. J. Stat. Softw. 32, 1–29 (2009).

89. Loytynoja, A. & Goldman, N. Phylogeny-aware gap placement prevents errors in sequence alignment and evolutionary analysis. Science 320, 1632–1635 (2008).

90. Ronquist, F. et al. MrBayes 3.2: efficient Bayesian phylogenetic inference and model choice across a large model space. Syst. Biol. 61, 539–542 (2012).

91. Szöllősi, G.J. et al. Lateral gene transfer from the dead. Syst. Biol. 62, 386–397 (2013).

92. Paradis, E. & Schliep, K. ape 5.0: an environment for modern phylogenetics and evolutionary analyses in R. Bioinformatics 35, 526–528 (2019).

93. Morris, J. L. et al. The timescale of early land plant evolution. Proc. Natl. Acad. Sci. U. S. A. 115, E2274–E2283 (2018).

94. Larsén, E. & Rydin, C. Disentangling the phylogeny of Isoetes (Isoetales), using nuclear and plastid data. Int. J. Plant Sci. 177, 157–174 (2016).

95. Edgar, R. C. MUSCLE: multiple sequence alignment with high accuracy and high throughput. Nucleic Acids Res. 32, 1792–1797 (2004).

96. Kalyaanamoorthy, S., Minh, B. Q., Wong, T. K. F., von Haeseler, A. & Jermiin, L. S. ModelFinder: fast model selection for accurate phylogenetic estimates. Nat. Methods 14, 587–589 (2017).

97. Nguyen, L.-T., Schmidt, H. A., von Haeseler, A. & Minh, B. Q. IQ-TREE: a fast and effective stochastic algorithm for estimating maximum-likelihood phylogenies. Mol. Biol. Evol. 32, 268–274 (2015).

98. Hoang, D. T., Chernomor, O., von Haeseler, A., Minh, B. Q. & Vinh, L. S. UFBoot2: Improving the Ultrafast Bootstrap Approximation. Mol. Biol. Evol. 35, 518–522 (2018).

99. de Vries, S. et al. The evolution of the phenylpropanoid pathway entailed pronounced radiations and divergences of enzyme families. bioRxiv 2021.05.27.445924 (2021) doi:10.1101/2021.05.27.445924.

100. Slotte, T. et al. The Capsella rubella genome and the genomic consequences of rapid mating system evolution. Nat. Genet. 45, 831–835 (2013).

101. Wan, T. et al. A genome for gnetophytes and early evolution of seed plants. Nat. Plants 4, 82–89 (2018).

102. Sierro, N. et al. The tobacco genome sequence and its comparison with those of tomato and potato. Nat. Commun. 5, 1–9 (2014).

103. Ouyang, S. et al. The TIGR Rice Genome Annotation Resource: improvements and new features. Nucleic Acids Res. 35, D883–7 (2007).

104. Argout, X. et al. The genome of Theobroma cacao. Nat. Genet. 43, 101–108 (2011).

105. Wang, S. et al. Genomes of early-diverging streptophyte algae shed light on plant terrestrialization. Nat Plants 6, 95–106 (2020).

106. Nishiyama, T. et al. The Chara genome: secondary complexity and implications for plant terrestrialization. Cell 174, 448–464.e24 (2018).

107. Hori, K. et al. Klebsormidium flaccidum genome reveals primary factors for plant terrestrial adaptation. Nat. Commun. 5, 3978 (2014).

108. Jiao, C. et al. The Penium margaritaceum genome: hallmarks of the origins of land plants. Cell 181, 1097–1111.e12 (2020).

109. de Vries, J. et al. Heat stress response in the closest algal relatives of land plants reveals conserved stress signaling circuits. Plant J. 103, 1025–1048 (2020).

110. de Vries, J., Curtis, B. A., Gould, S. B. & Archibald, J. M. Embryophyte stress signaling evolved in the algal progenitors of land plants. Proc. Natl. Acad. Sci. U. S. A. 115, E3471– E3480 (2018).

111. Ju, C. et al. Conservation of ethylene as a plant hormone over 450 million years of evolution. Nat. Plants 1, 14004 (2015).

112. Moreau, H. et al. Gene functionalities and genome structure in Bathycoccus prasinos reflect cellular specializations at the base of the green lineage. Genome Biol. 13, R74 (2012).

113. Merchant, S. S. et al. The Chlamydomonas genome reveals the evolution of key animal and plant functions. Science 318, 245–250 (2007).

114. Blanc, G. et al. The genome of the polar eukaryotic microalga Coccomyxa subellipsoidea reveals traits of cold adaptation. Genome Biol. 13, R39 (2012).

115. Worden, A. Z. et al. Green evolution and dynamic adaptations revealed by genomes of the marine picoeukaryotes Micromonas. Science 324, 268–272 (2009).

116. Palenik, B. et al. The tiny eukaryote Ostreococcus provides genomic insights into the paradox of plankton speciation. Proc. Natl. Acad. Sci. U. S. A. 104, 7705–7710 (2007).

117. De Clerck, O. et al. Insights into the evolution of multicellularity from the sea lettuce genome. Curr. Biol. 28, 2921–2933.e5 (2018).

118. Prochnik, S. E. et al. Genomic analysis of organismal complexity in the multicellular green alga Volvox carteri. Science 329, 223–226 (2010).

119. de Vries, J., de Vries, S., Slamovits, C. H., Rose, L. E. & Archibald, J. M. How embryophytic is the biosynthesis of phenylpropanoids and their derivatives in Streptophyte algae? Plant Cell Physiol. 58, 934–945 (2017).

120. Katoh, K. & Standley, D. M. MAFFT multiple sequence alignment software version 7: improvements in performance and usability. Mol. Biol. Evol. 30, 772–780 (2013).

121. Minh, B. Q. et al. IQ-TREE 2: New models and efficient methods for phylogenetic inference in the genomic Era. Mol. Biol. Evol. 37, 1530–1534 (2020).

122. Hu, Y. et al. Crystal structures of a Populus tomentosa 4-coumarate:CoA ligase shed light on its enzymatic mechanisms. Plant Cell 22, 3093–3104 (2010).

123. Pan, H. et al. Structural studies of cinnamoyl-CoA reductase and cinnamyl-alcohol dehydrogenase, key enzymes of monolignol biosynthesis. Plant Cell 26, 3709–3727 (2014).

124. Louie, G. V. et al. Structure-function analyses of a caffeic acid O-methyltransferase from perennial ryegrass reveal the molecular basis for substrate preference. Plant Cell 22, 4114–4127 (2010).

125. Youn, B. et al. Crystal structures and catalytic mechanism of the Arabidopsis cinnamyl alcohol dehydrogenases AtCAD5 and AtCAD4. Org. Biomol. Chem. 4, 1687–1697 (2006).

126. Ferrer, J.-L., Zubieta, C., Dixon, R. A. & Noel, J. P. Crystal structures of alfalfa caffeoyl coenzyme A 3-O-methyltransferase. Plant Physiol. 137, 1009–1017 (2005).

127. Barbier, F. F. et al. A phenol/chloroform-free method to extract nucleic acids from recalcitrant, woody tropical species for gene expression and sequencing. Plant Methods 15, 62 (2019).

128. Love, M., Anders, S. & Huber, W. Differential analysis of count data--the DESeq2 package. Genome Biol. 15, 10–1186 (2014).

129. Conesa, A., Nueda, M. J., Ferrer, A. & Talón, M. maSigPro: a method to identify significantly differential expression profiles in time-course microarray experiments. Bioinformatics 22, 1096–1102 (2006).

130. MacKinnon, Kirk J-M., et al. Changes in ambient temperature are the prevailing cue in determining Brachypodium distachyon diurnal gene regulation. New Phytologist 227, 1709–1724 (2020).

131. Michael, T. P. et al. A morning-specific phytohormone gene expression program underlying rhythmic plant growth. PLoS Biol. 6, e225 (2008).

132. Mockler, T. C. et al. The DIURNAL project: DIURNAL and circadian expression profiling, model-based pattern matching, and promoter analysis. Cold Spring Harb. Symp. Quant. Biol. 72, 353–363 (2007).

133. McLeay, R. C. & Bailey, T. L. Motif Enrichment Analysis: a unified framework and an evaluation on ChIP data. BMC Bioinformatics 11, 165 (2010).

134. Franco-Zorrilla, J. M. et al. DNA-binding specificities of plant transcription factors and their potential to define target genes. Proc. Natl. Acad. Sci. U. S. A. 111, 2367–2372 (2014).

135. Grant, C. E., Bailey, T. L. & Noble, W. S. FIMO: scanning for occurrences of a given motif. Bioinformatics 27, 1017–1018 (2011).

