## Supplementary Figures for "Underwater CAM photosynthesis elucidated by *Isoetes* genome"

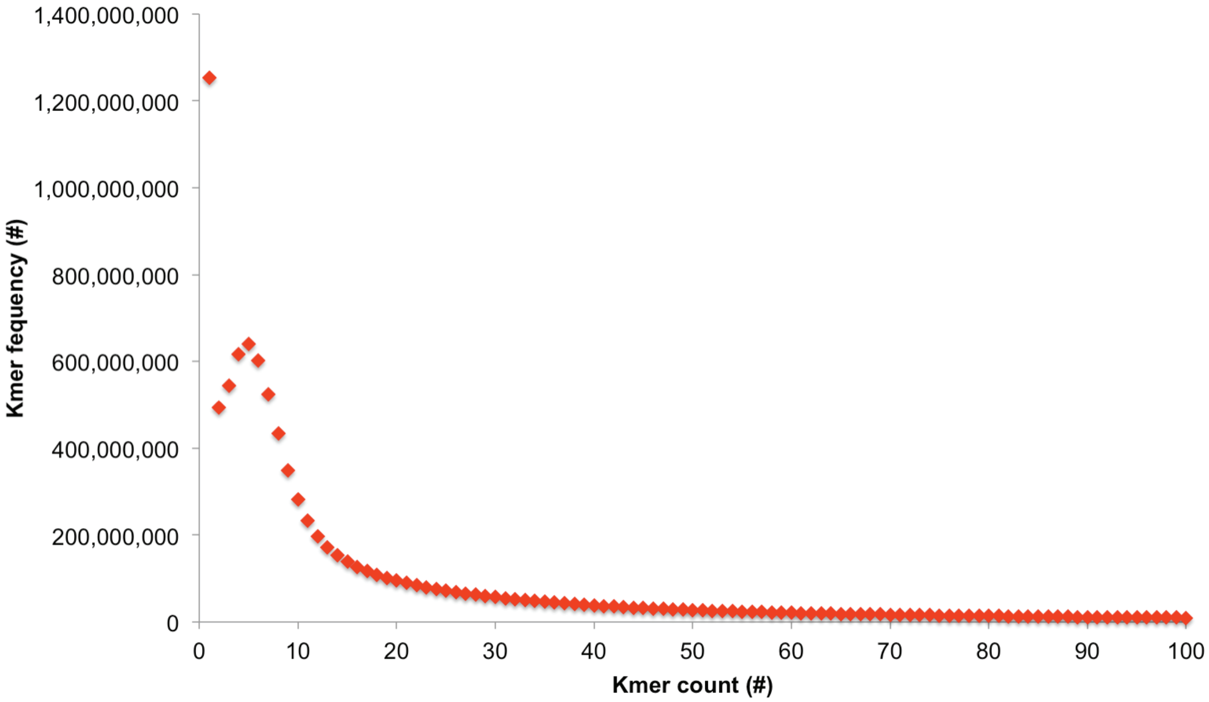


**Supplementary Figure S1: *Isoetes taiwanensis* genome size estimate based on short read k-mer frequency.** The genome size was estimated to be 1,647,045,703 bp by k-mer frequency (k=19) analysis using Illumina 2x150 bp short reads.

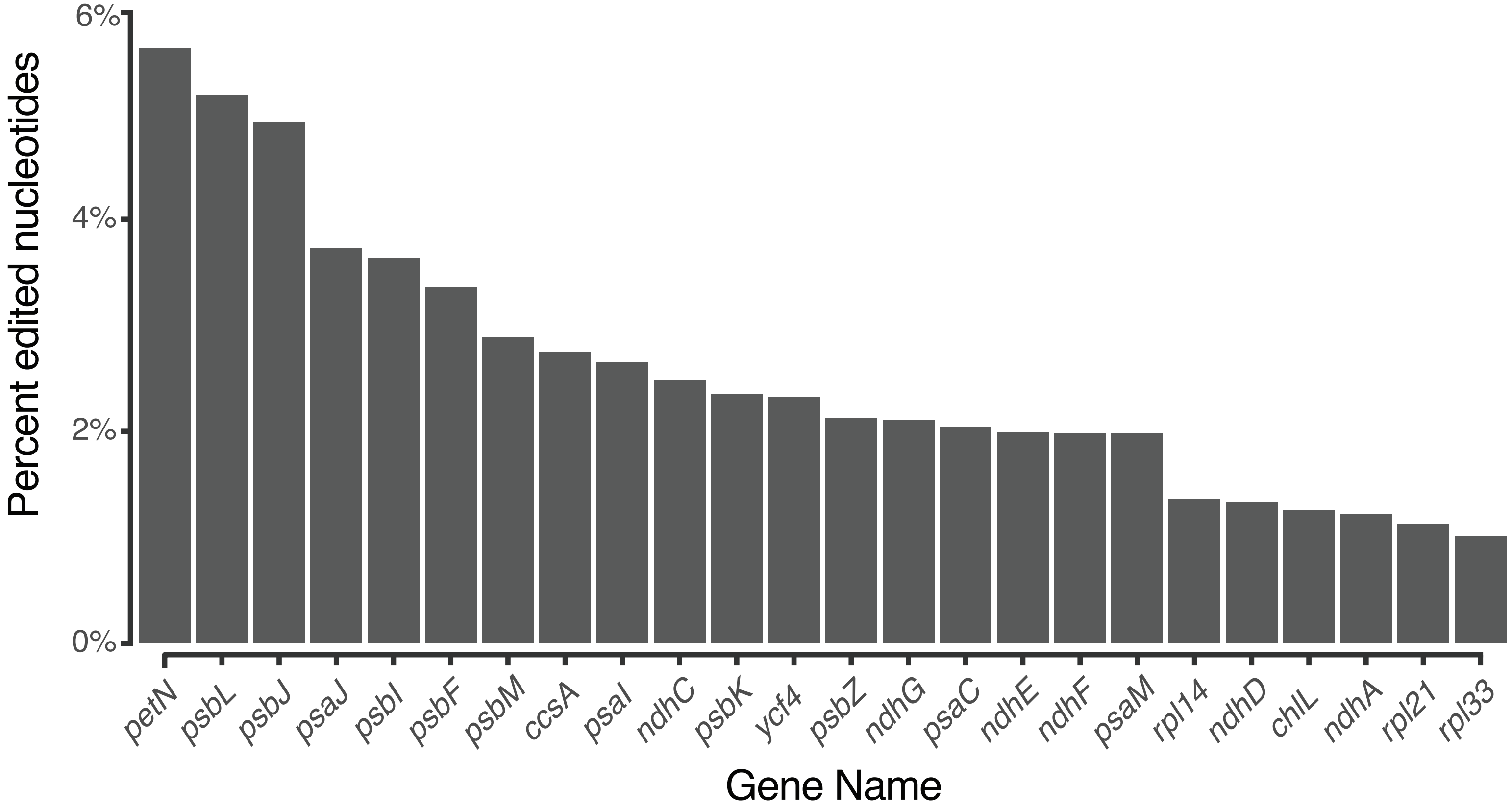


**Supplemental Figure S2: The *Isoetes taiwanensis* plastome shows a high degree of RNA editing.** A plot showing the number of editing sites as a percentage of gene length in genes with greater than 1% edited nucleotides.

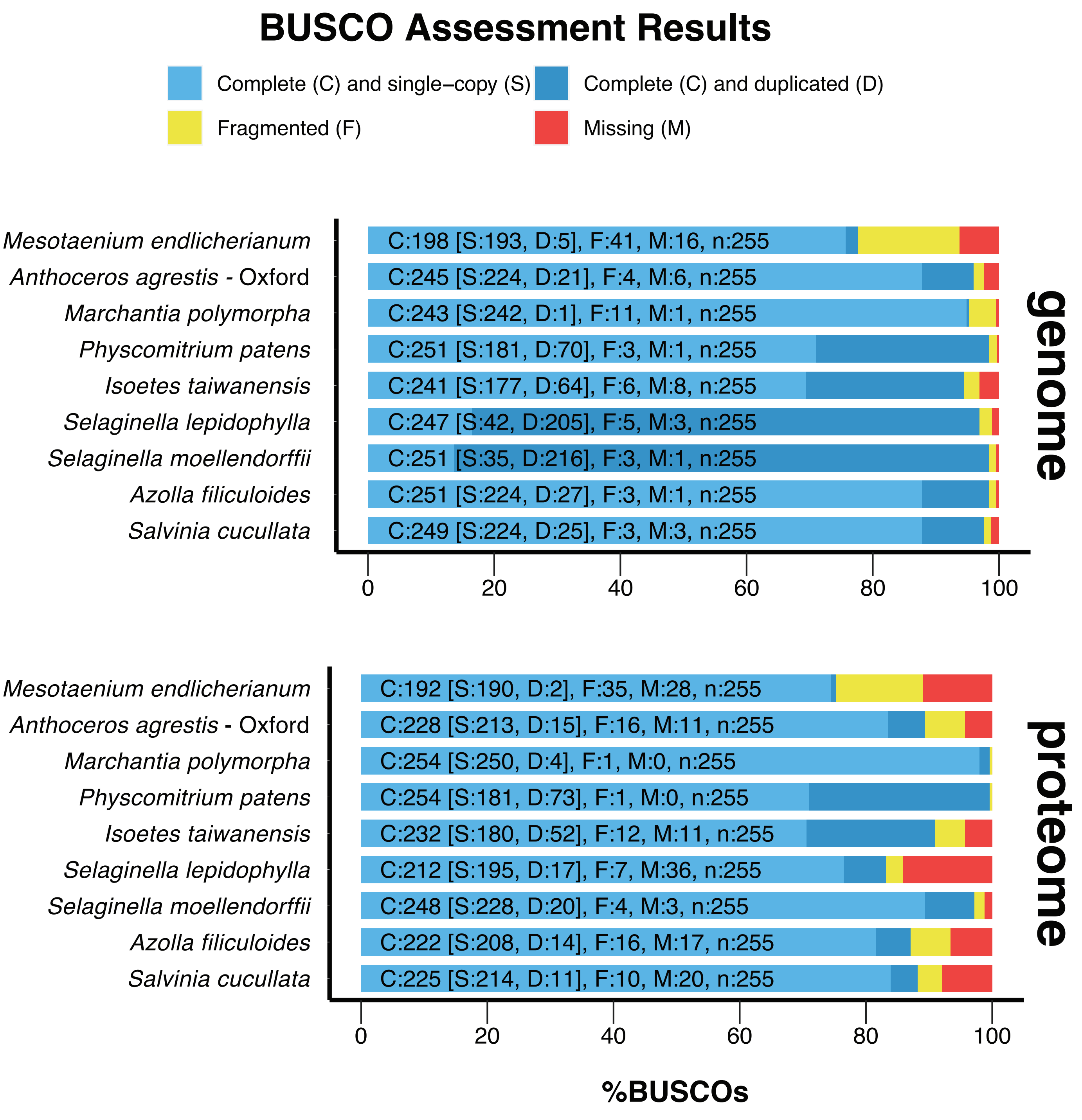


**Supplemental Figure S3: Comparison of BUSCO scores for *Isoetes taiwanensis* and other seed-free plant genomes.** BUSCO analysis showed that the *I. taiwanensis* genome assembly is comparable to other seed-free plant and charophyte genome assemblies with regard to completeness.

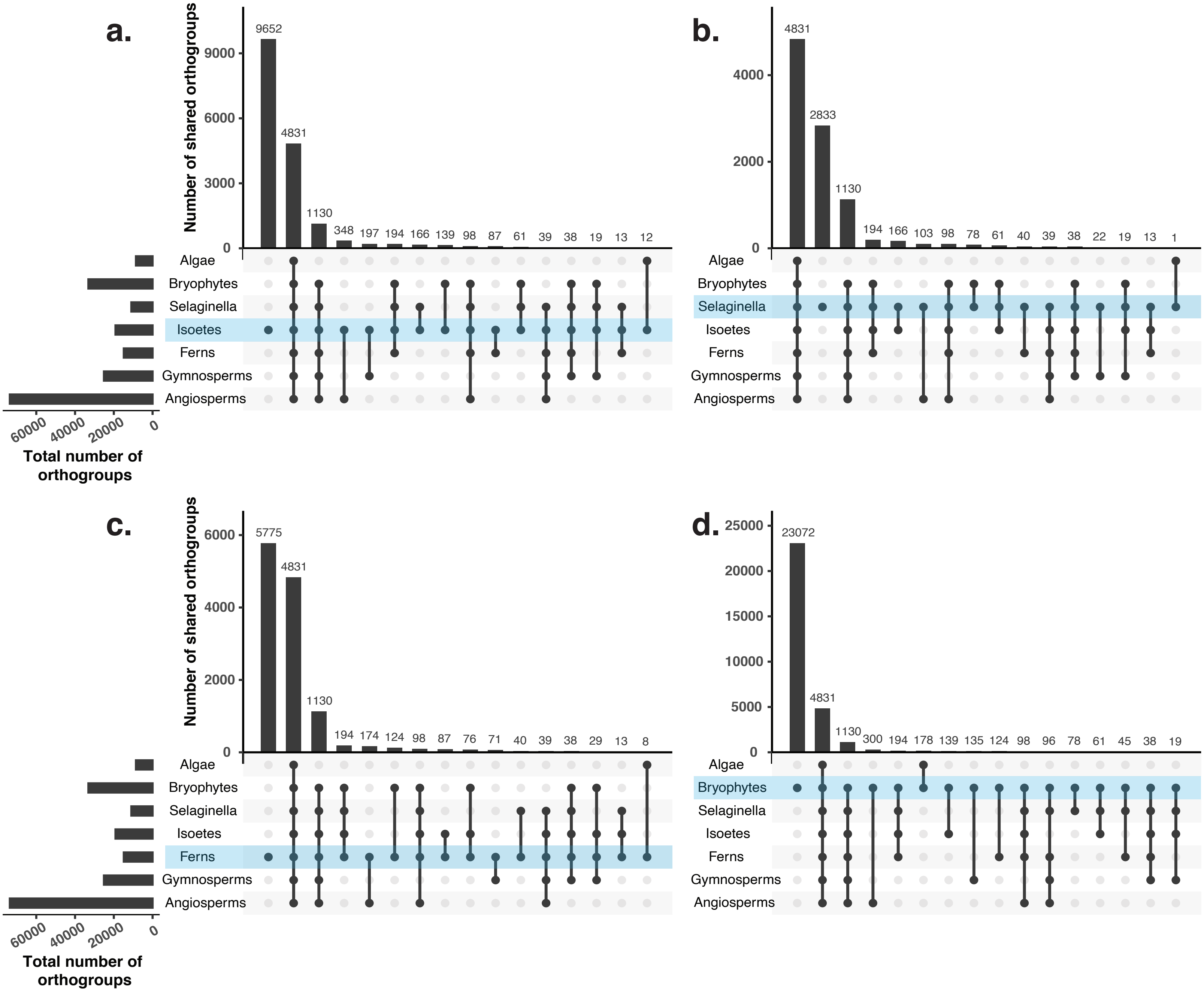


**Supplemental Figure S4: Orthogroups in *Isoetes* have greater overlap with seed plants than seed-free taxa.** Upset plots show degree of overlap among various plant lineages included in Orthofinder analysis. Each plot focuses on shared orthogroups with a particular group: **a**, *Isoetes taiwanensis*, **b**, *Selaginella moellendorffii*, **c**, ferns, and **d**, bryophytes.

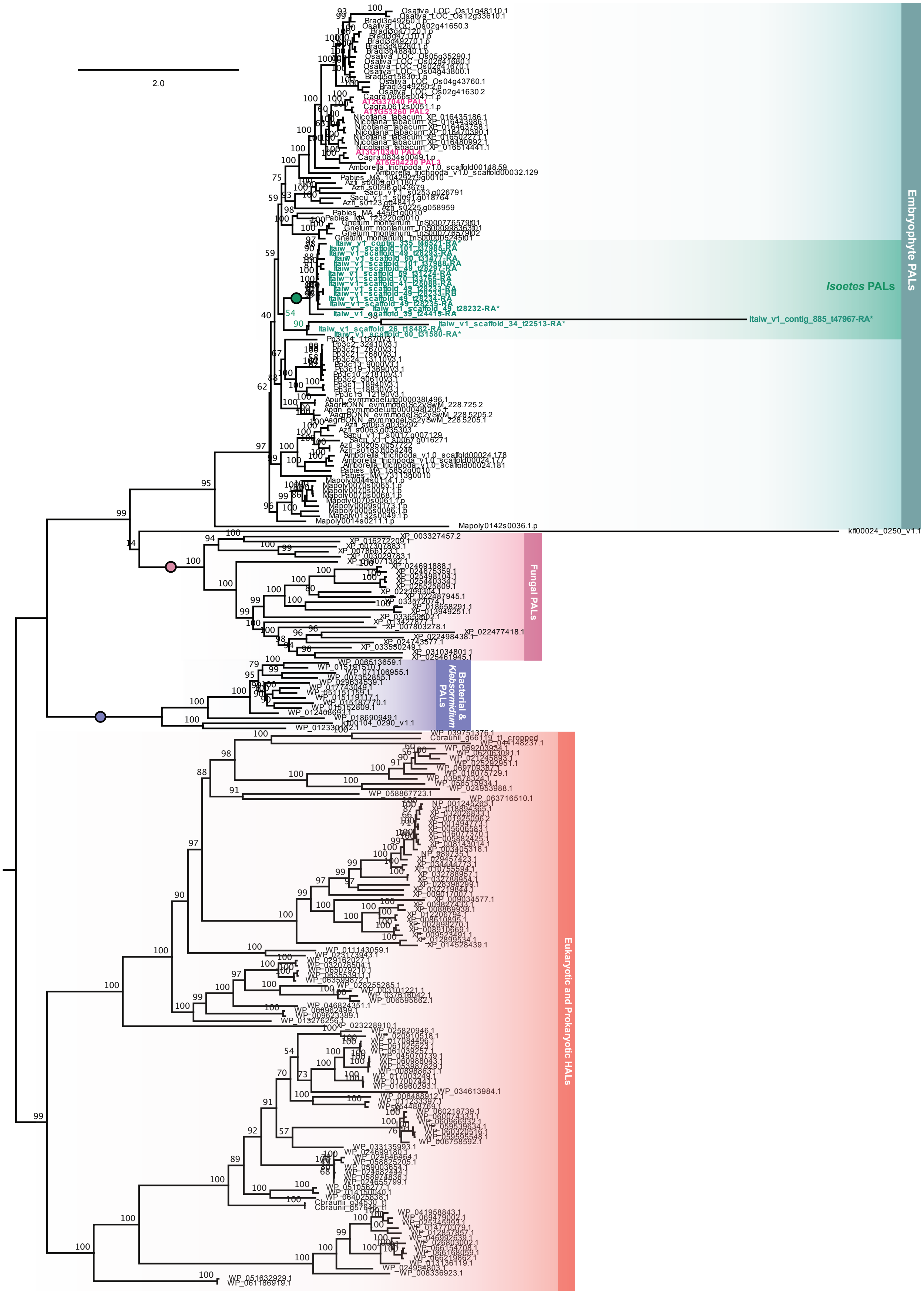


**Supplemental Figure S5: Rooted phylogeny of phenylalanine ammonia lyase (PAL) homologs in *Isoetes taiwanensis* and across Chloroplastida.** Full bootstrap support for highlighted clades is indicated by a filled circle. Unusually short *I. taiwanensis* sequences in highlighted clades are labeled with an asterisk.

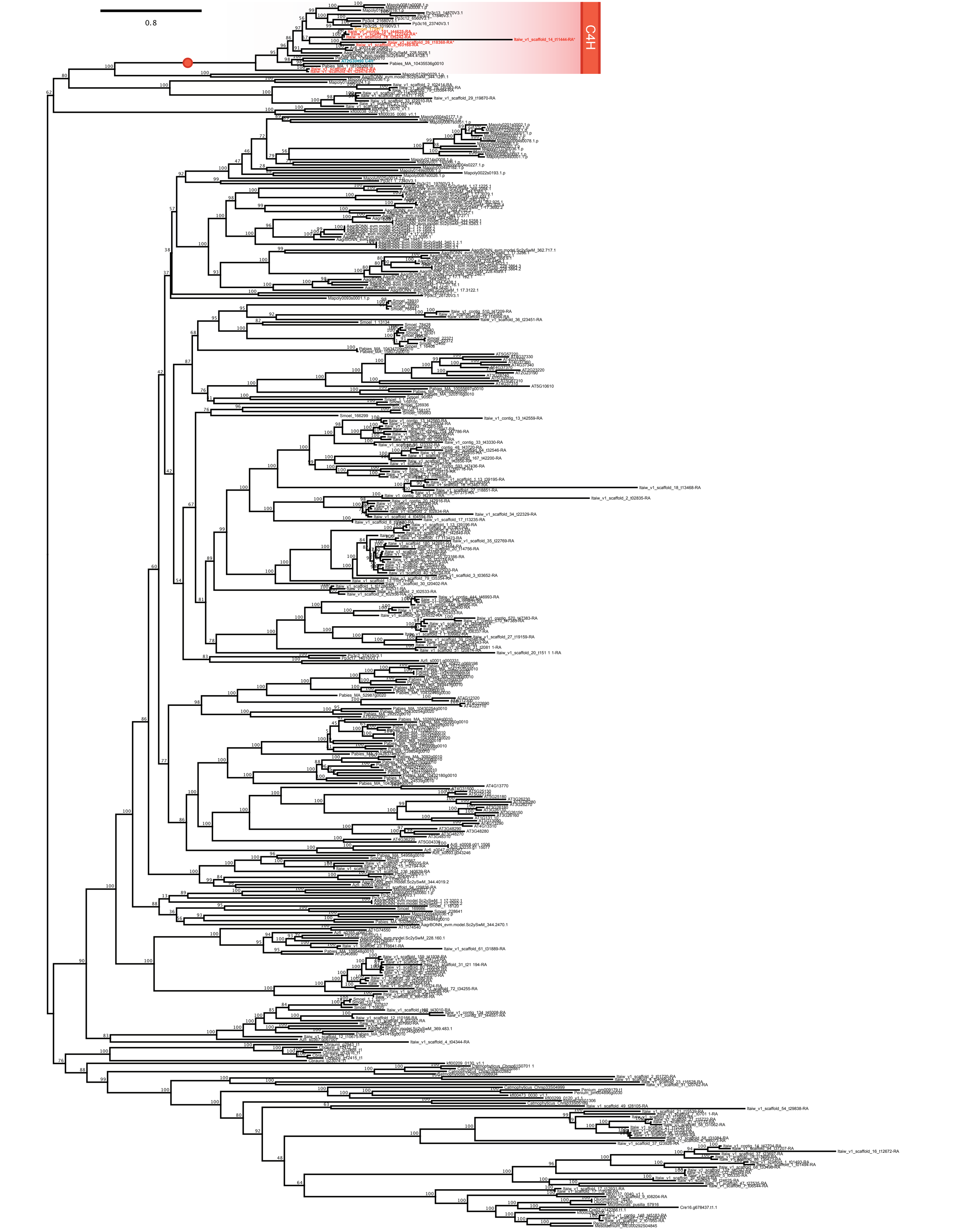


**Supplemental Figure S6: Unrooted phylogeny of cinnamate-4-hydroxylase (C4H) homologs in *Isoetes taiwanensis* and across Chloroplastida*.*** Relevant clades are labeled using gradient color; homologues of note are highlighted with colored font (teal for *Arabidopsis*, orange for *Selaginella moellendorffii*, red for *I. taiwanensis*). Full bootstrap support for highlighted clades is indicated by a filled circle. Unusually short *I. taiwanensis* sequences in highlighted clades are labeled with an asterisk.

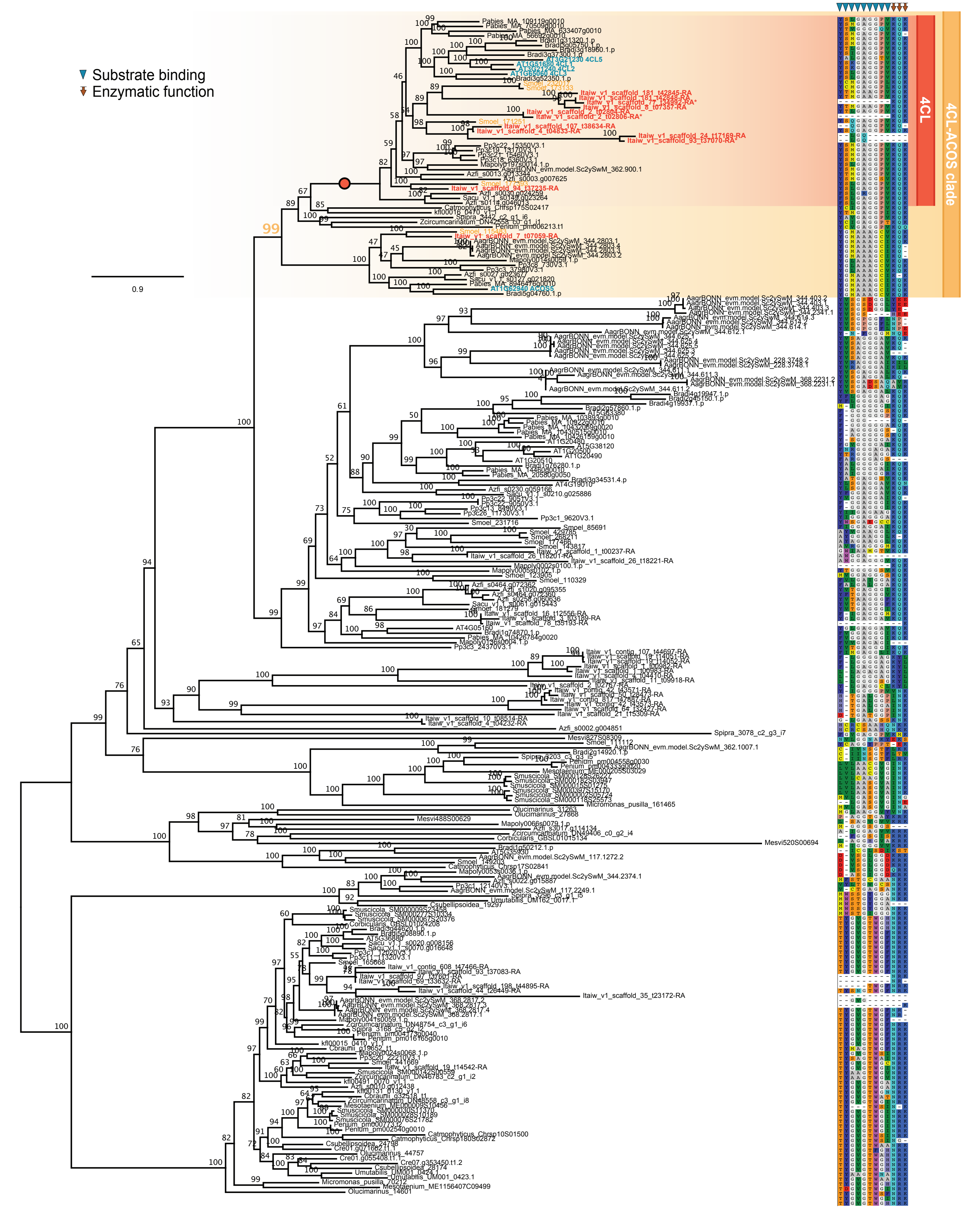


**Supplemental Figure S7: Unrooted phylogeny of 4-coumarate CoA ligase (4CL) homologs in *Isoetes taiwanensis* and across Chloroplastida.** Relevant clades are labeled using gradient color; homologues of note are highlighted with colored font (teal for *Arabidopsis*, orange for *Selaginella moellendorffii*, red for *I. taiwanensis*). Full bootstrap support for highlighted clades is indicated by a filled circle. To the right of the phylogeny residues relevant to substrate binding and function of canonical 4CL as reported by Hu et al.[1] are shown. Unusually short *I. taiwanensis* sequences in highlighted clades are labeled with an asterisk.

*1. Hu, Y. et al. Crystal structures of a Populus tomentosa 4-coumarate:CoA ligase shed light on its enzymatic mechanisms. Plant Cell 22, 3093–3104 (2010).*

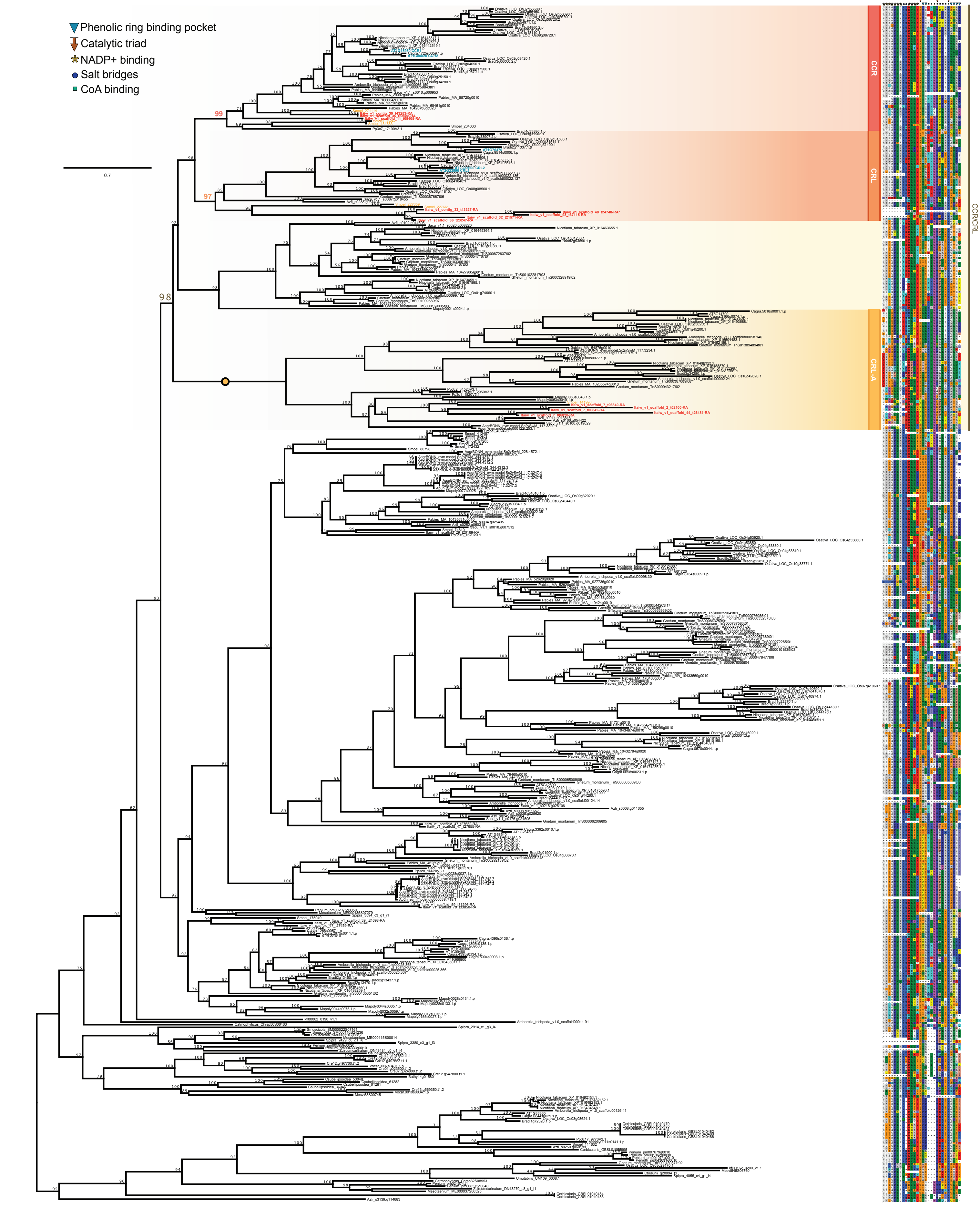


**Supplemental Figure S8: Unrooted phylogeny of cinnamoyl CoA reductase (CCR) homologs in *Isoetes taiwanensis* and across Chloroplastida.** Relevant clades are labeled using gradient color; homologues of note are highlighted with colored font (teal for *Arabidopsis*, orange for *Selaginella moellendorffii*, red for *I. taiwanensis*). Full bootstrap support for highlighted clades is indicated by a filled circle. To the right of the phylogeny residues relevant to phenolic ring binding pocket, the catalytic triad, NADP+ binding, salt bridges, and CoA binding of CCR as reported by Pan et al. [1] are shown. Unusually short *I. taiwanensis* sequences in highlighted clades are labeled with an asterisk.

*1.Pan, H. et al. Structural studies of cinnamoyl-CoA reductase and cinnamyl-alcohol dehydrogenase, key enzymes of monolignol biosynthesis. Plant Cell 26, 3709–3727 (2014).*

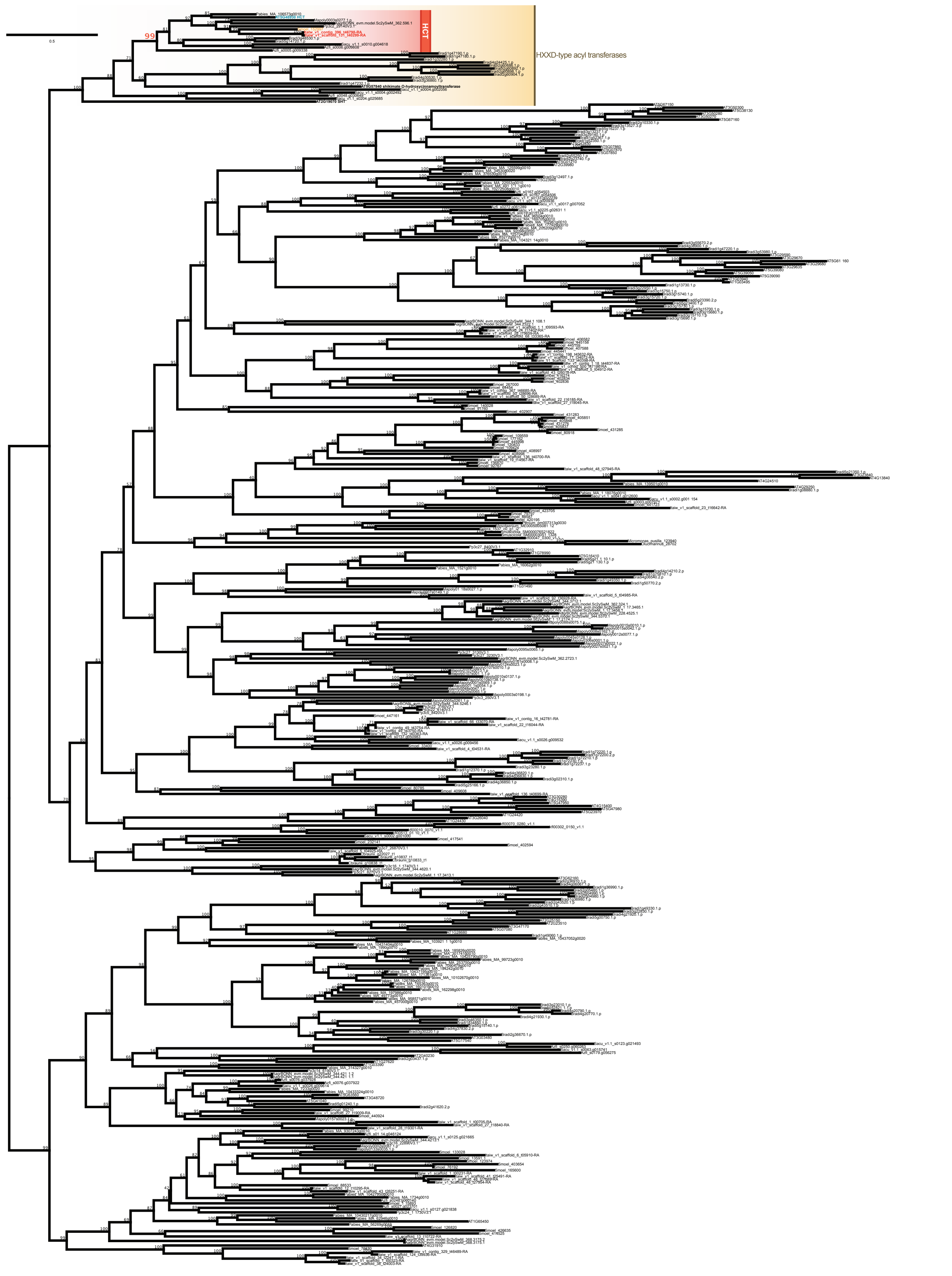


**Supplemental Figure S9: Unrooted phylogeny of hydroxycinnamoyl-Coenzyme A shikimae/quinate hydroxycinnamoyl-transferase (HCT) homologs in *Isoetes taiwanensis* and across Chloroplastida.** Relevant clades are labeled using gradient color; homologues of note are highlighted with colored font (teal for *Arabidopsis*, orange for *Selaginella moellendorffii*, red for *I. taiwanensis*). Full bootstrap support for highlighted clades is indicated by a filled circle.

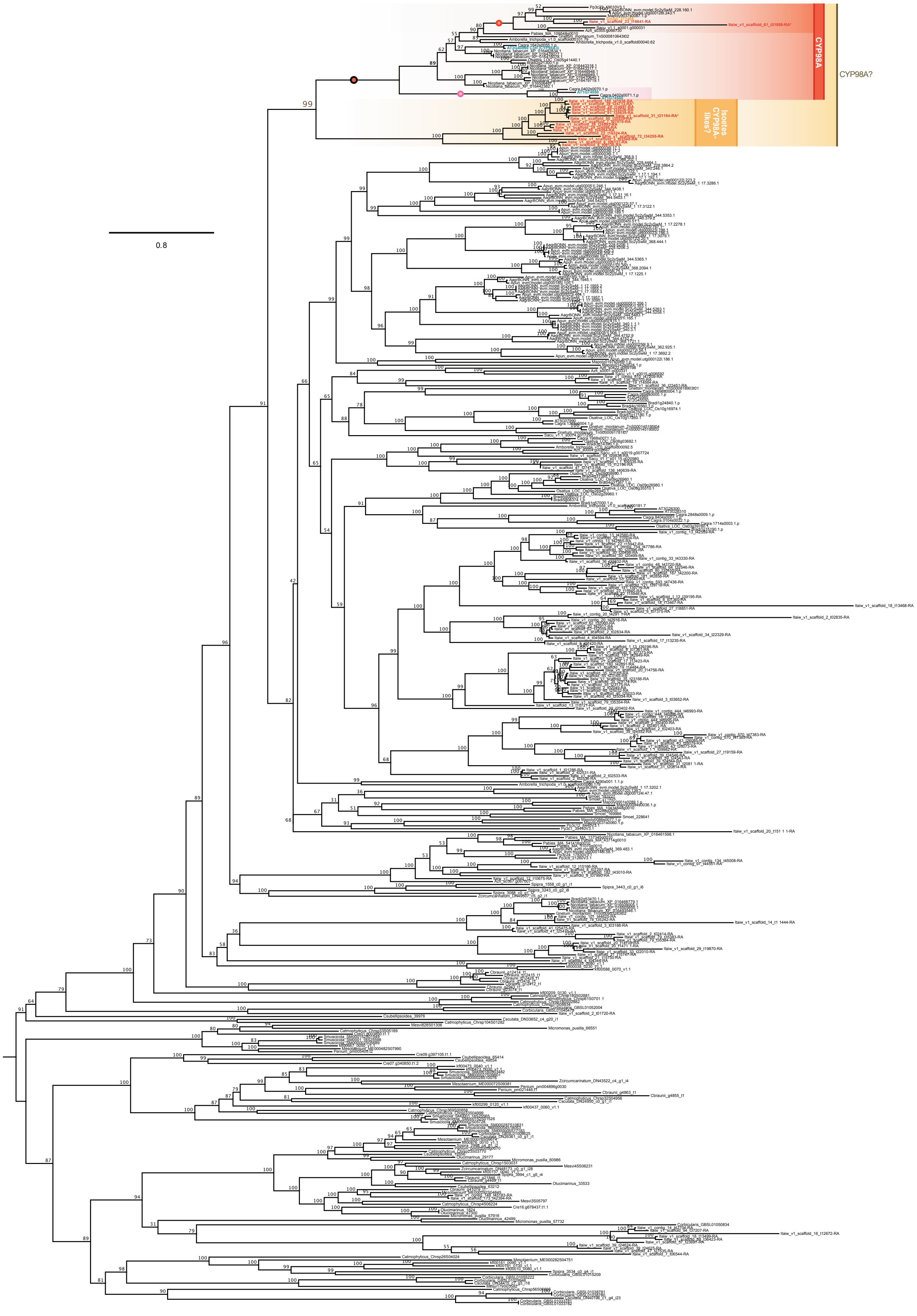


**Supplemental Figure S10: Unrooted phylogeny of coumarate 3-hydroxylase (C3H) homologs in *Isoetes taiwanensis* and across Chloroplastida.** Relevant clades are labeled using gradient color; homologues of note are highlighted with colored font (teal for *Arabidopsis*, orange for *Selaginella moellendorffii*, red for *I. taiwanensis*). Full bootstrap support for highlighted clades is indicated by a filled circle. Unusually short *I. taiwanensis* sequences in highlighted clades are labeled with an asterisk.

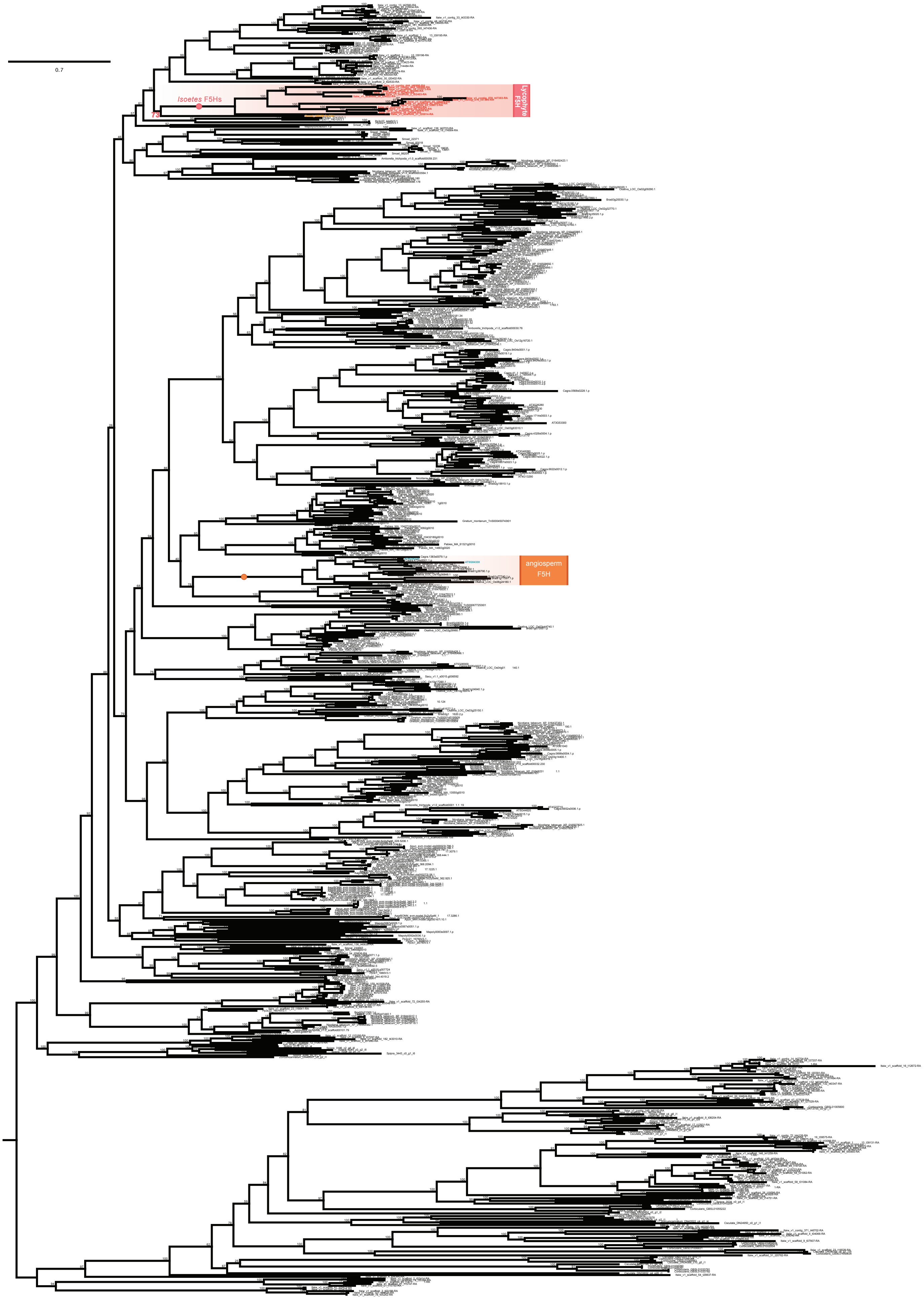


**Supplemental Figure S11: Unrooted phylogeny of ferulate 5-hydroxylase (F5H) homologs in *Isoetes taiwanensis* and across Chloroplastida*.*** Relevant clades are labeled using gradient color; homologues of note are highlighted with colored font (teal for *Arabidopsis*, orange for *Selaginella moellendorffii*, red for *I. taiwanensis*). Full bootstrap support for highlighted clades is indicated by a filled circle.

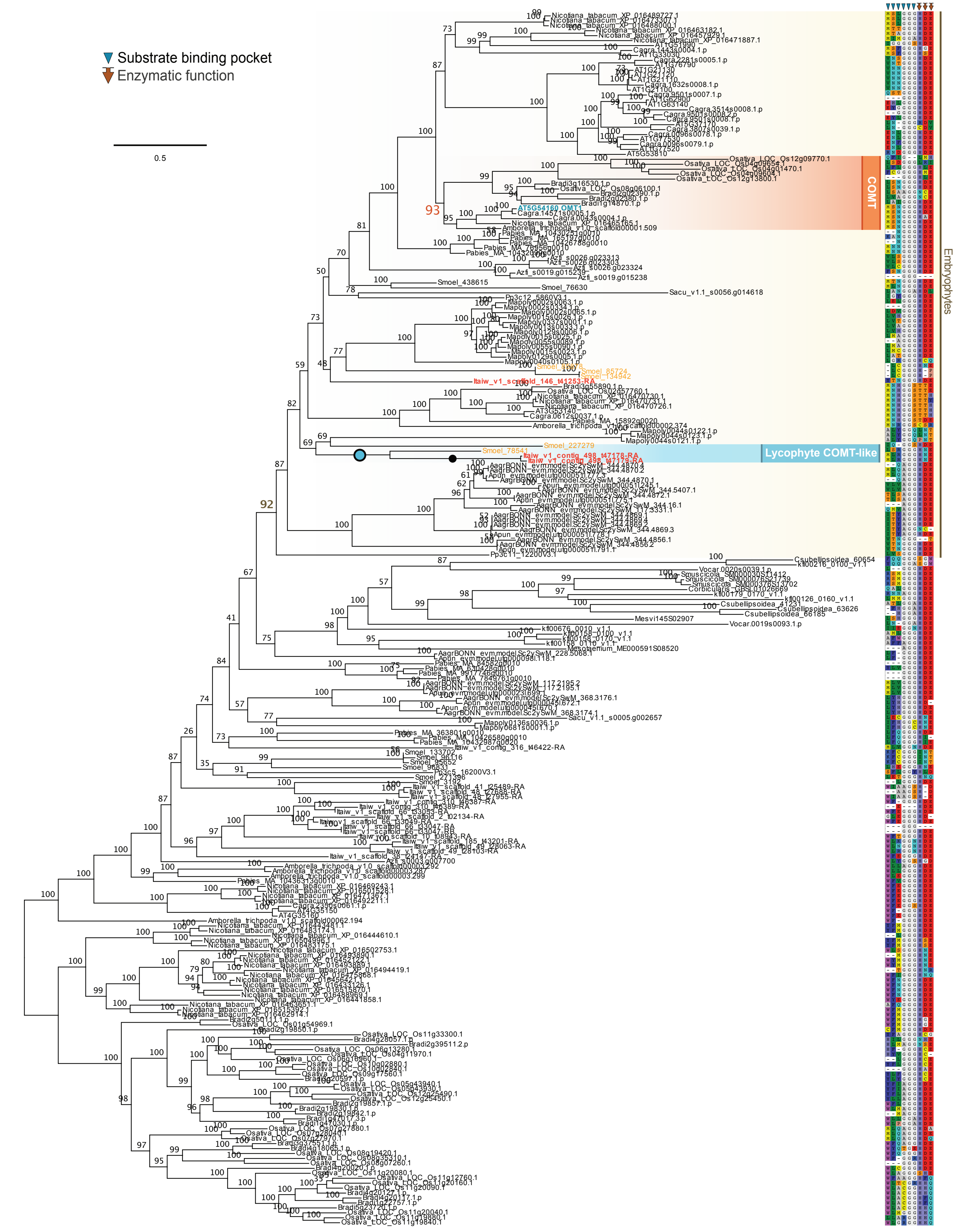


**Supplemental Figure S12: Unrooted phylogeny of caffeic acid/5-hydroxyferulic acid O-methyltransferase (COMT) homologs in *Isoetes taiwanensis* and across Chloroplastida.** Relevant clades are labeled using gradient color; homologues of note are highlighted with colored font (teal for *Arabidopsis*, orange for *Selaginella moellendorffii*, red for *I. taiwanensis*). Full bootstrap support for highlighted clades is indicated by a filled circle. To the right of the phylogeny residues relevant to substrate binding and function of COMT as reported by Louie et al.[1] are shown.

*1. Louie, G. V. et al. Structure-function analyses of a caffeic acid O-methyltransferase from perennial ryegrass reveal the molecular basis for substrate preference. Plant Cell 22, 4114–4127 (2010).*

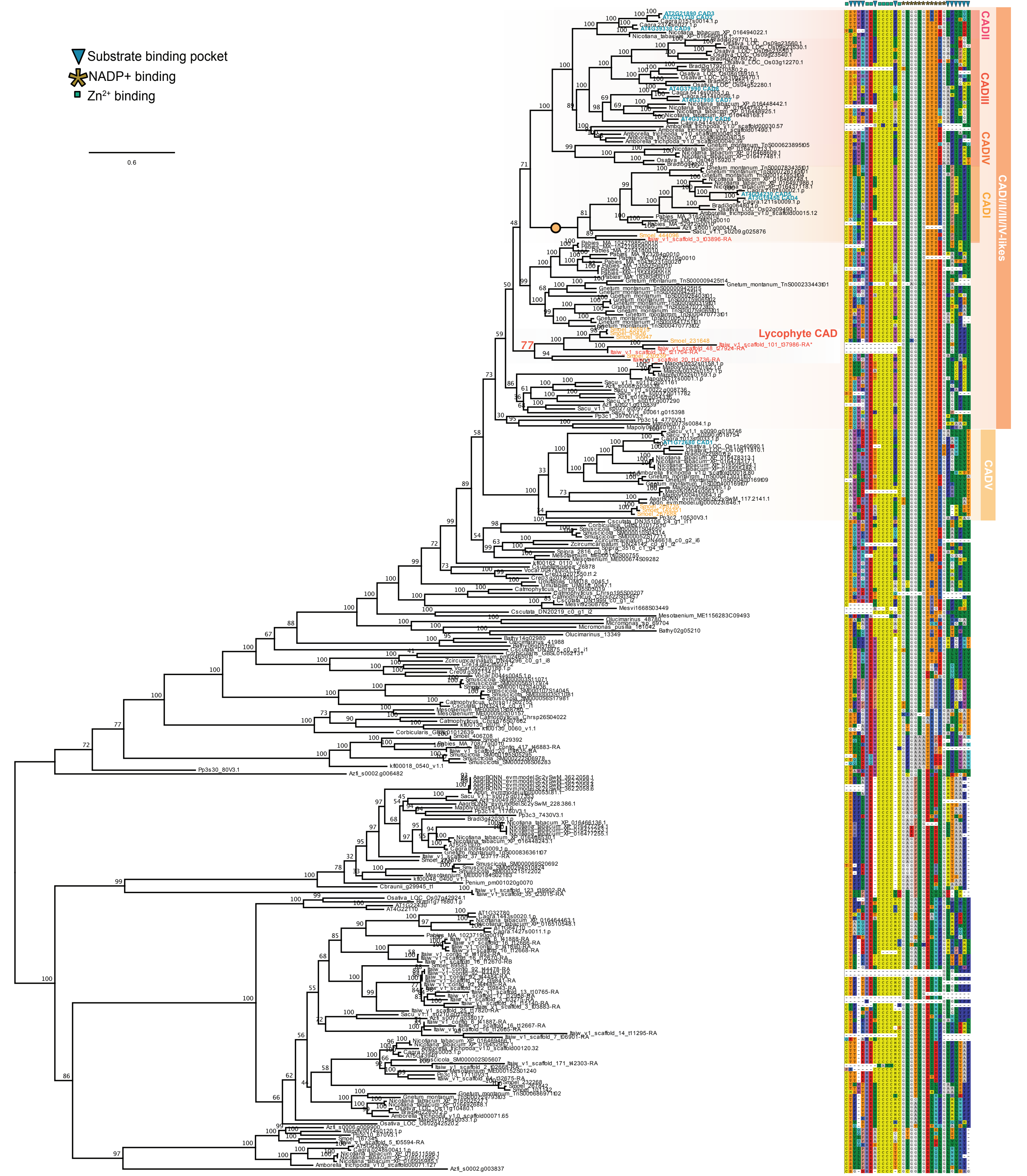


**Supplemental Figure S13: Unrooted phylogeny of cinnamyl-alcohol dehydrogenase (CAD) homologs in *Isoetes taiwanensis* and across Chloroplastida.** Relevant clades are labeled using gradient color; homologues of note are highlighted with colored font (teal for *Arabidopsis*, orange for *Selaginella moellendorffii*, red for *I. taiwanensis*). Full bootstrap support for highlighted clades is indicated by a filled circle. To the right of the phylogeny residues relevant to substrate binding and function of CAD binding pocket, NADP+- and $Zn^{2+}$ binding according to Youn et al.[1] as reported by Louie et al.[2] are shown. Unusually short *I. taiwanensis* sequences in highlighted clades are labeled with an asterisk.

*1. Youn, B. et al. Crystal structures and catalytic mechanism of the Arabidopsis cinnamyl alcohol dehydrogenases AtCAD5 and AtCAD4. Org. Biomol. Chem. 4, 1687–1697 (2006).*

*2. Louie, G. V. et al. Structure-function analyses of a caffeic acid O-methyltransferase from perennial ryegrass reveal the molecular basis for substrate preference. Plant Cell 22, 4114–4127 (2010).*

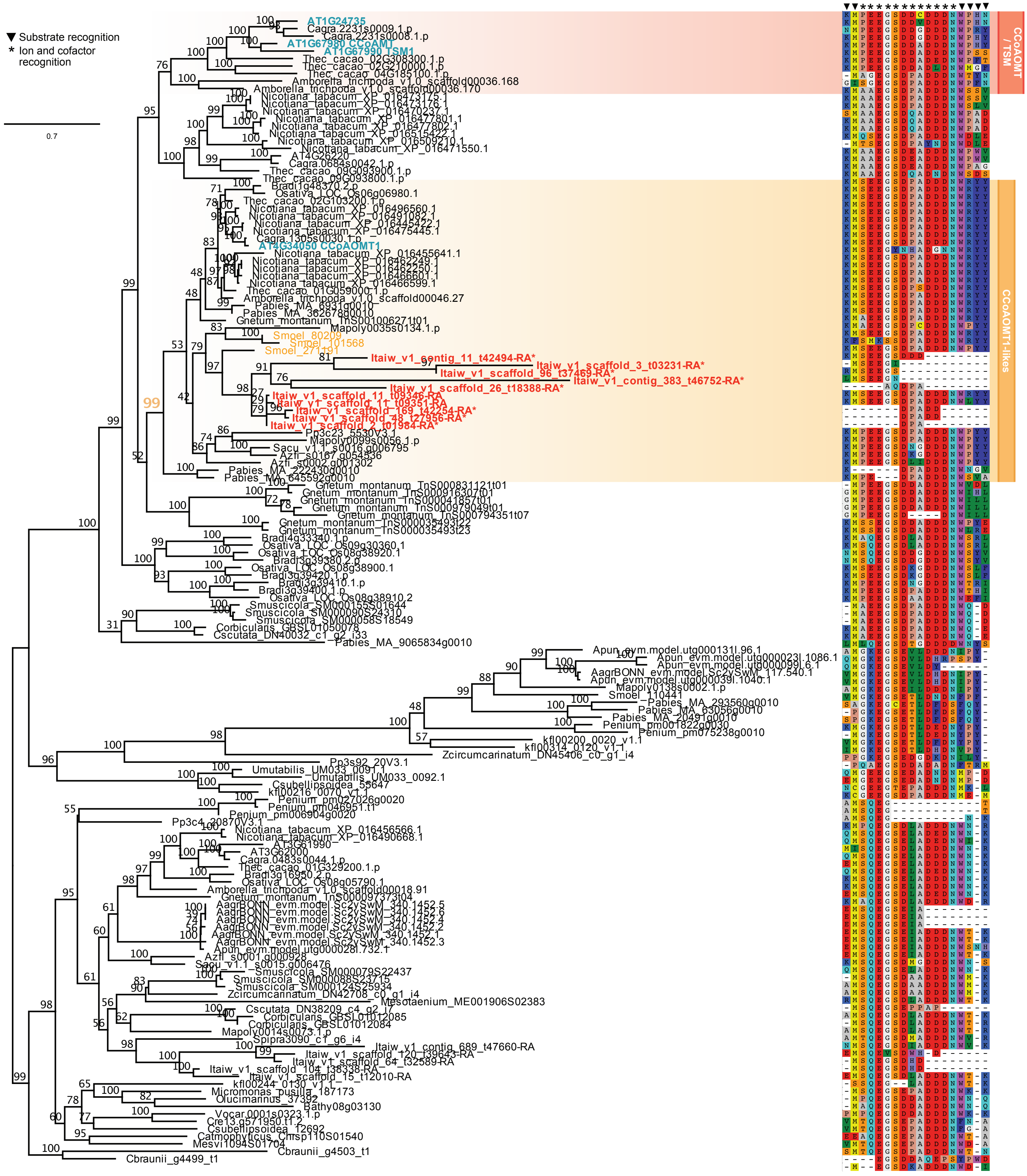


**Supplemental Figure S14: Unrooted phylogeny of caffeoyl-CoA O-methyltransferase (CCoAOMT) homologs in *Isoetes taiwanensis* and across Chloroplastida.** Relevant clades are labeled using gradient color; homologues of note are highlighted with colored font (teal for *Arabidopsis*, orange for *Selaginella moellendorffii*, red for *I. taiwanensis*). Full bootstrap support for highlighted clades is indicated by a filled circle. To the right of the phylogeny residues relevant to substrate, ion and cofactor recognition by CCoAOMT as reported by Ferrer et al.[1] are shown. Unusually short *I. taiwanensis* sequences in highlighted clades are labeled with an asterisk.

*1. Ferrer, J.-L., Zubieta, C., Dixon, R. A. & Noel, J. P. Crystal structures of alfalfa caffeoyl coenzyme A 3-O-methyltransferase. Plant Physiol. 137, 1009–1017 (2005).*

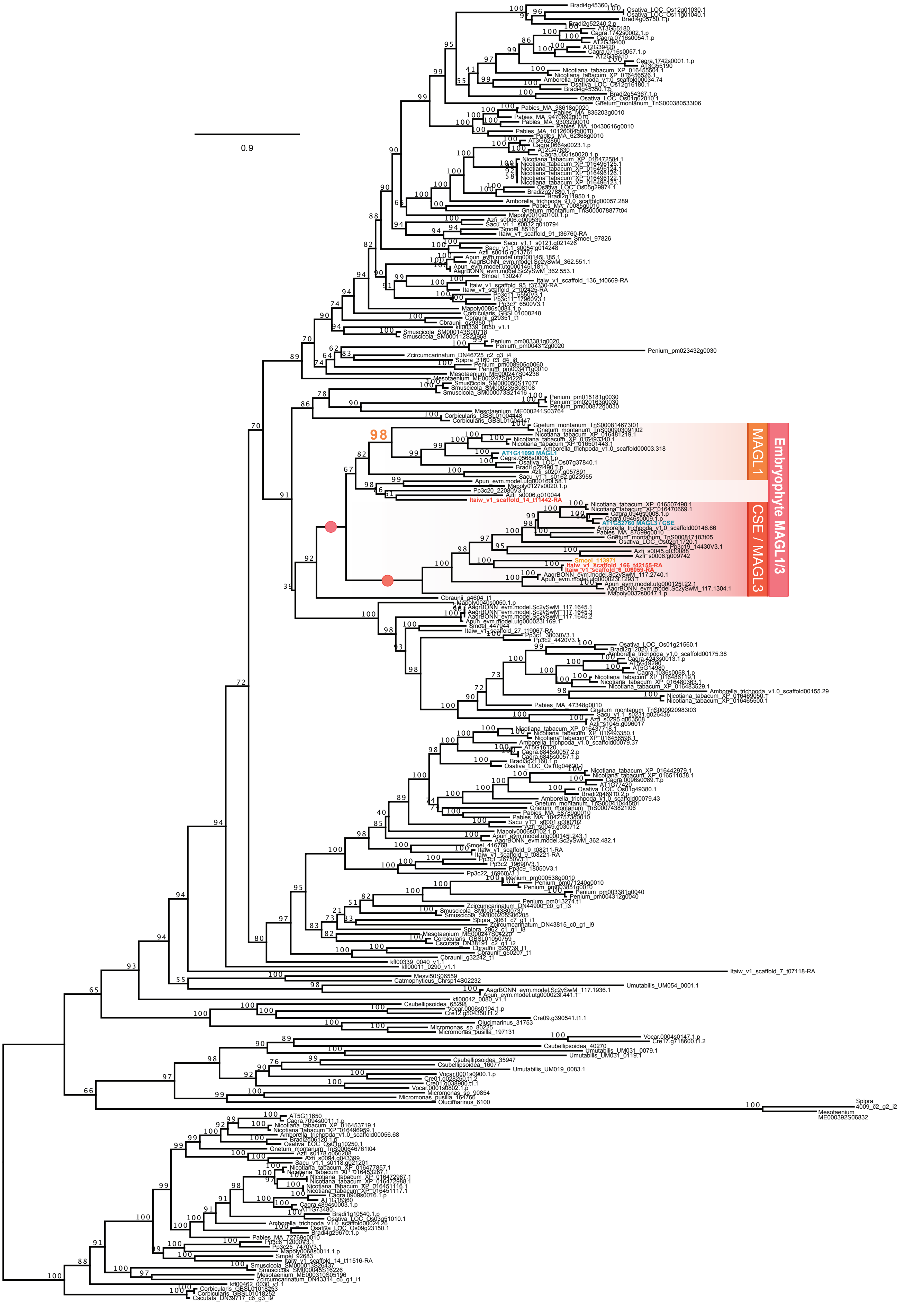


**Supplemental Figure S15: Unrooted phylogeny of caffeoyl shikimate esterase (CSE) / monoacylglycerol lipase (MAGL) homologs in *Isoetes taiwanensis* and across Chloroplastida.** Relevant clades are labeled using gradient color; homologues of note are highlighted with colored font (teal for *Arabidopsis*, orange for *Selaginella moellendorffii*, red for *I. taiwanensis*). Full bootstrap support for highlighted clades is indicated by a filled circle.

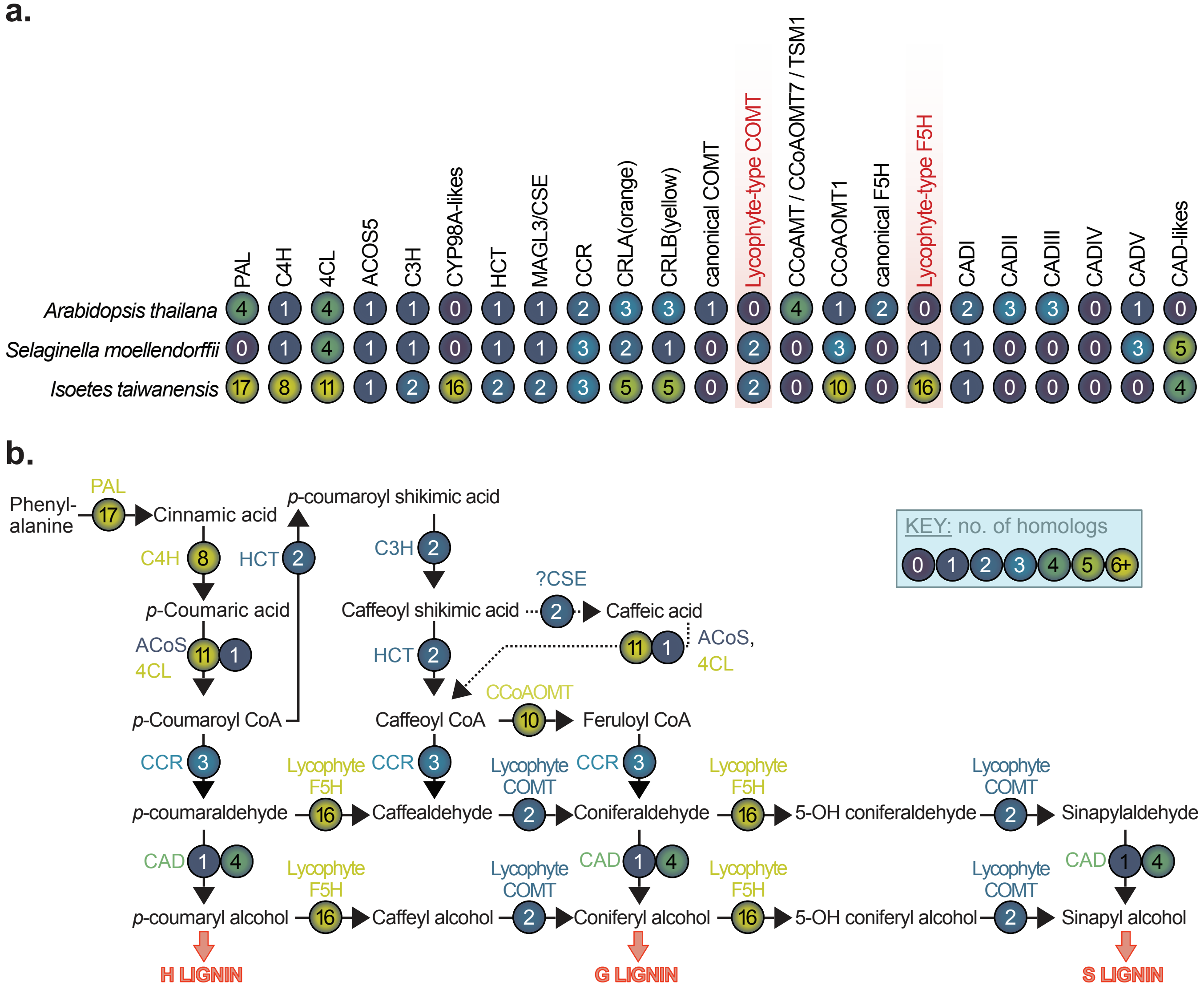


**Supplemental Figure S16: Overview of the detected homologues for key enzymes of the phenylpropanoid and lignin biosynthesis pathway in *Isoetes taiwanensis.* a,** Phylogenetically detected homologs for key enzymes of the core phenylpropanoid pathway as well as routes towards H, G, and S lignin are distributed across 23 groups. **b,** Putative and simplified biosynthetic routes toward lignin in *I. taiwanensis* as inferred from homologs detected. In both **a** and **b**, the number of detected homologs are shown in gradient colored bubbles.

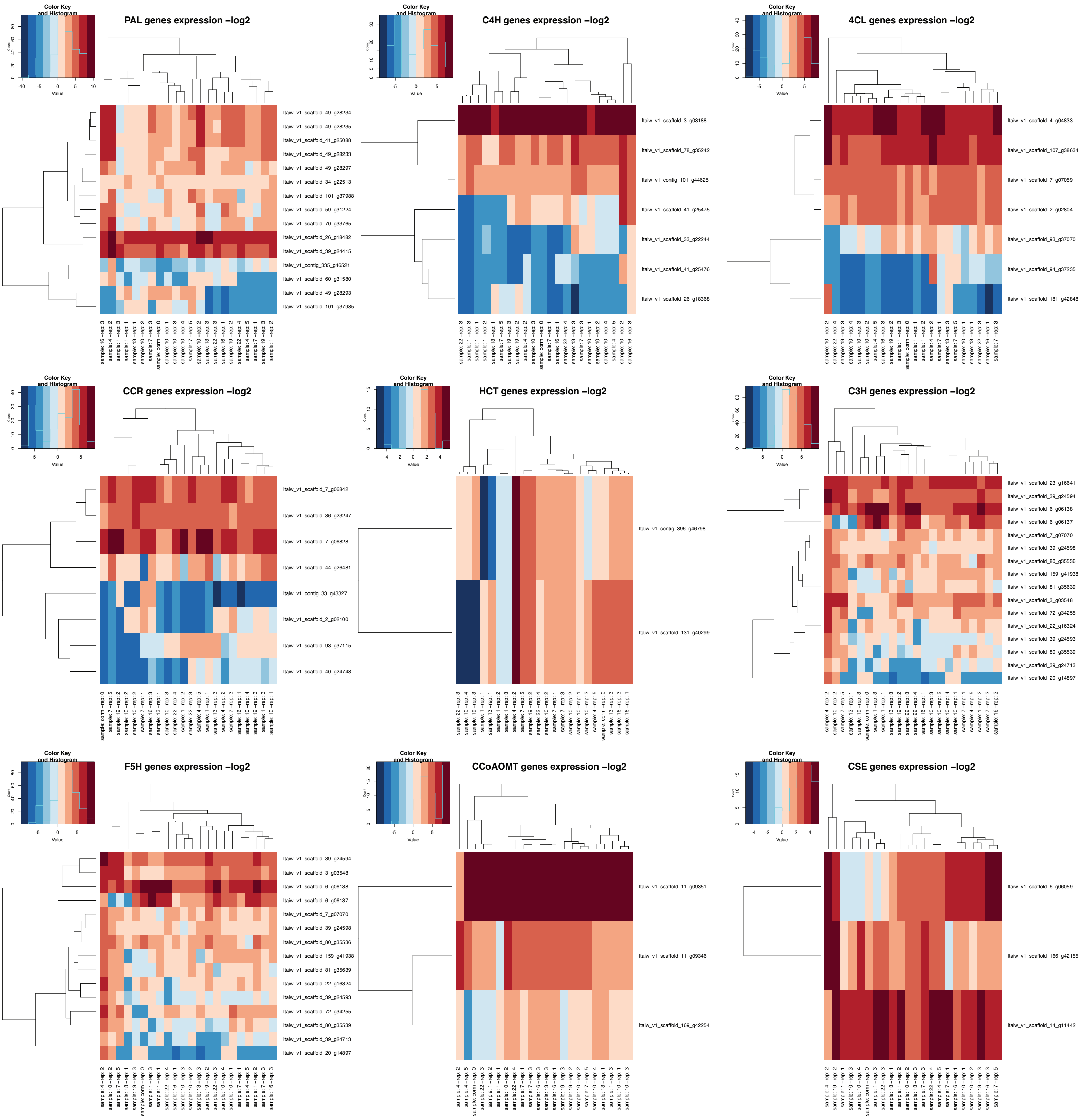


**Supplemental Figure S17: Expression of genes in the phenylpropanoid and lignin biosynthesis pathway in *Isoetes taiwanensis* .** Note that no COMT homologues and only a single copy of CAD were expressed. The latter is labeled in the phylogeny in Supplemental Figure S13.

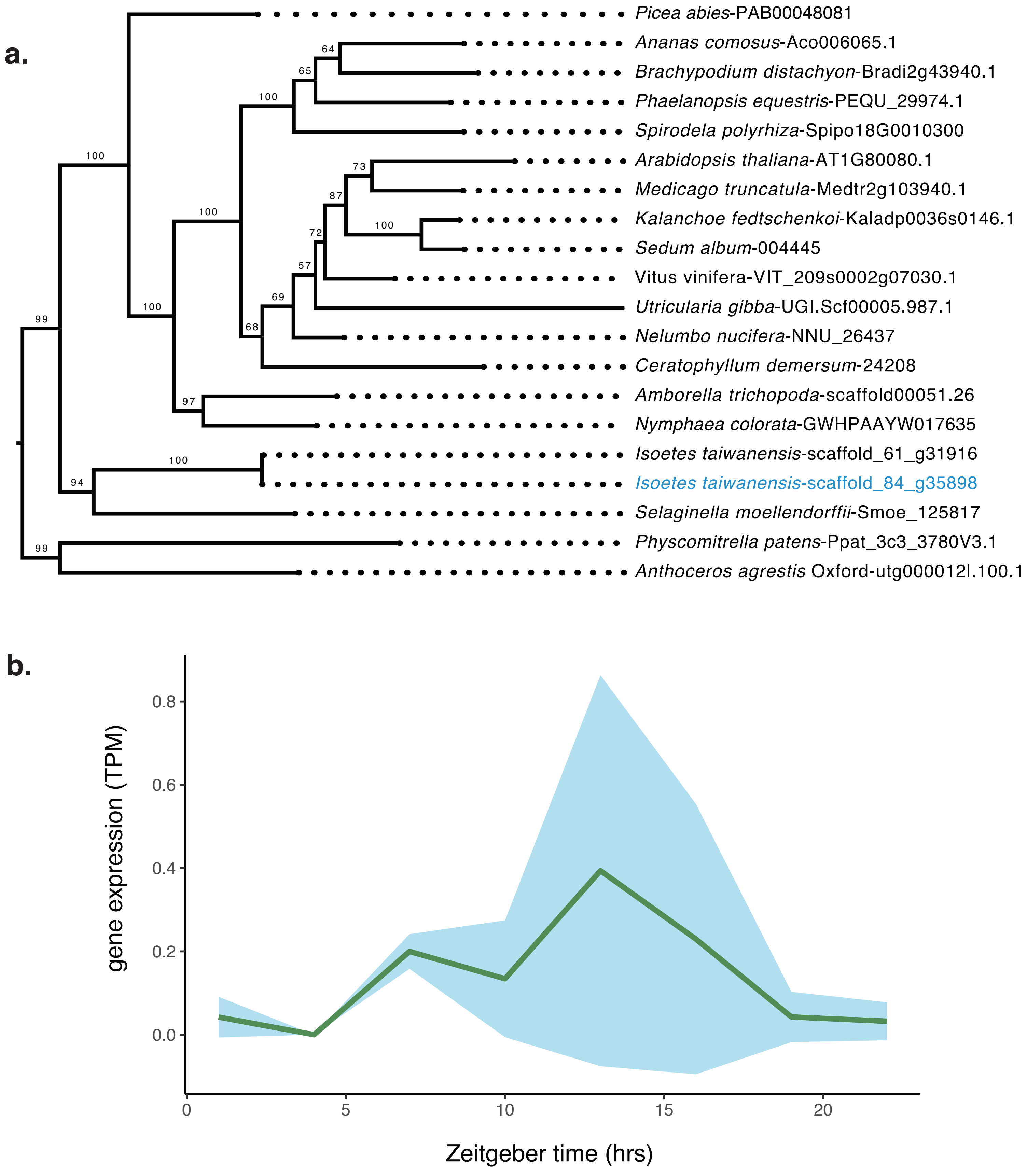


**Supplementary Figure S18: Orthologues of TMM found in *Isoetes taiwanensis*. a,** Phylogenetic placement of two copies of the stomatal development gene TMM from *I. taiwanensis* with **b,** normalized expression data from one copy expressed in leaf tissue. The copy of TMM expressed in leaf tissue is highlighted using blue text in the phylogeny.

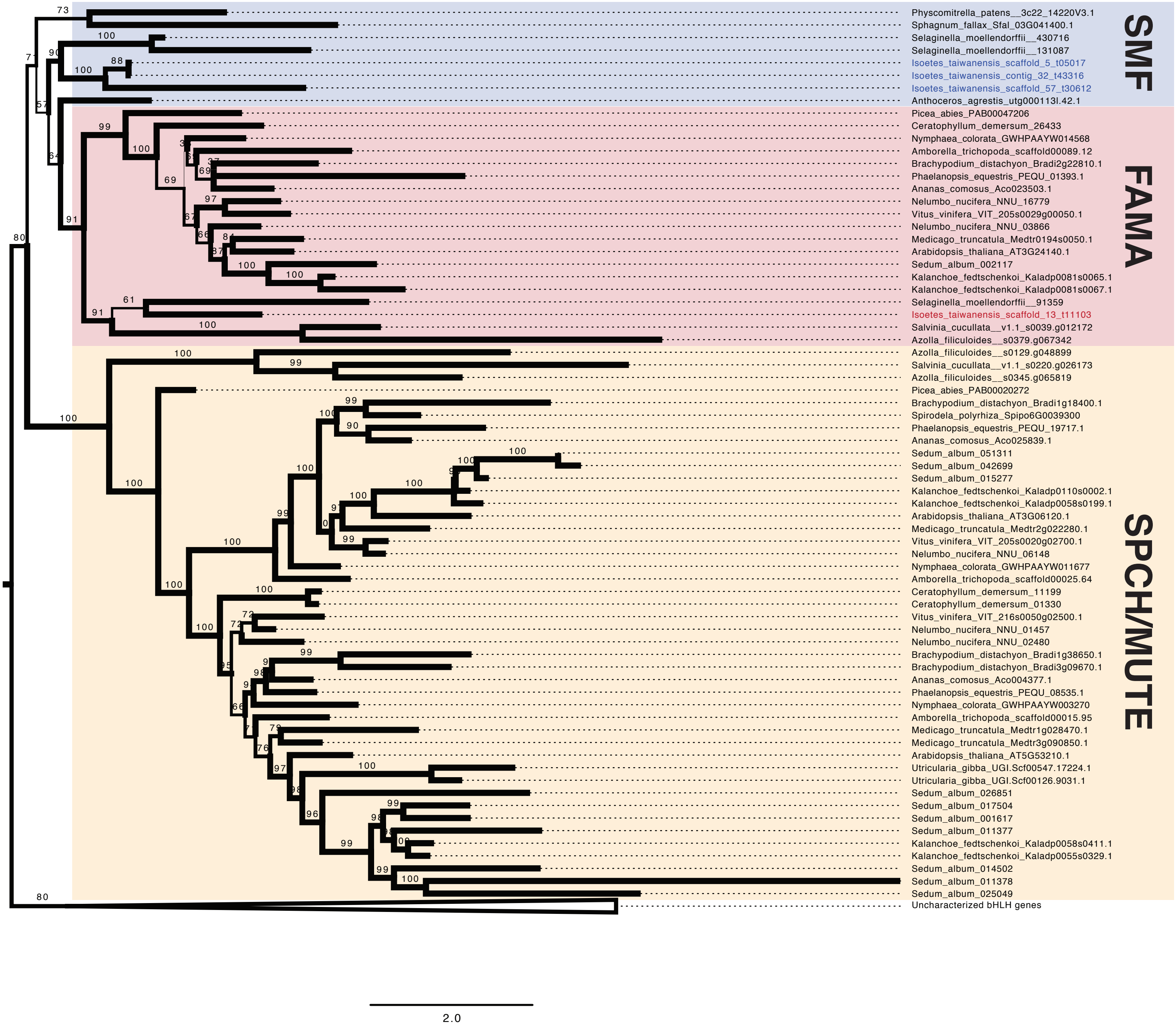


**Supplemental Figure S19: *Isoetes taiwanensis* possesses stomatal development genes in the SMF/FAMA clade.** A phylogeny showing the placement of SMF and FAMA genes found in *I. taiwanensis* relative to other land plants. Branch thickness corresponds to bootstrap support.

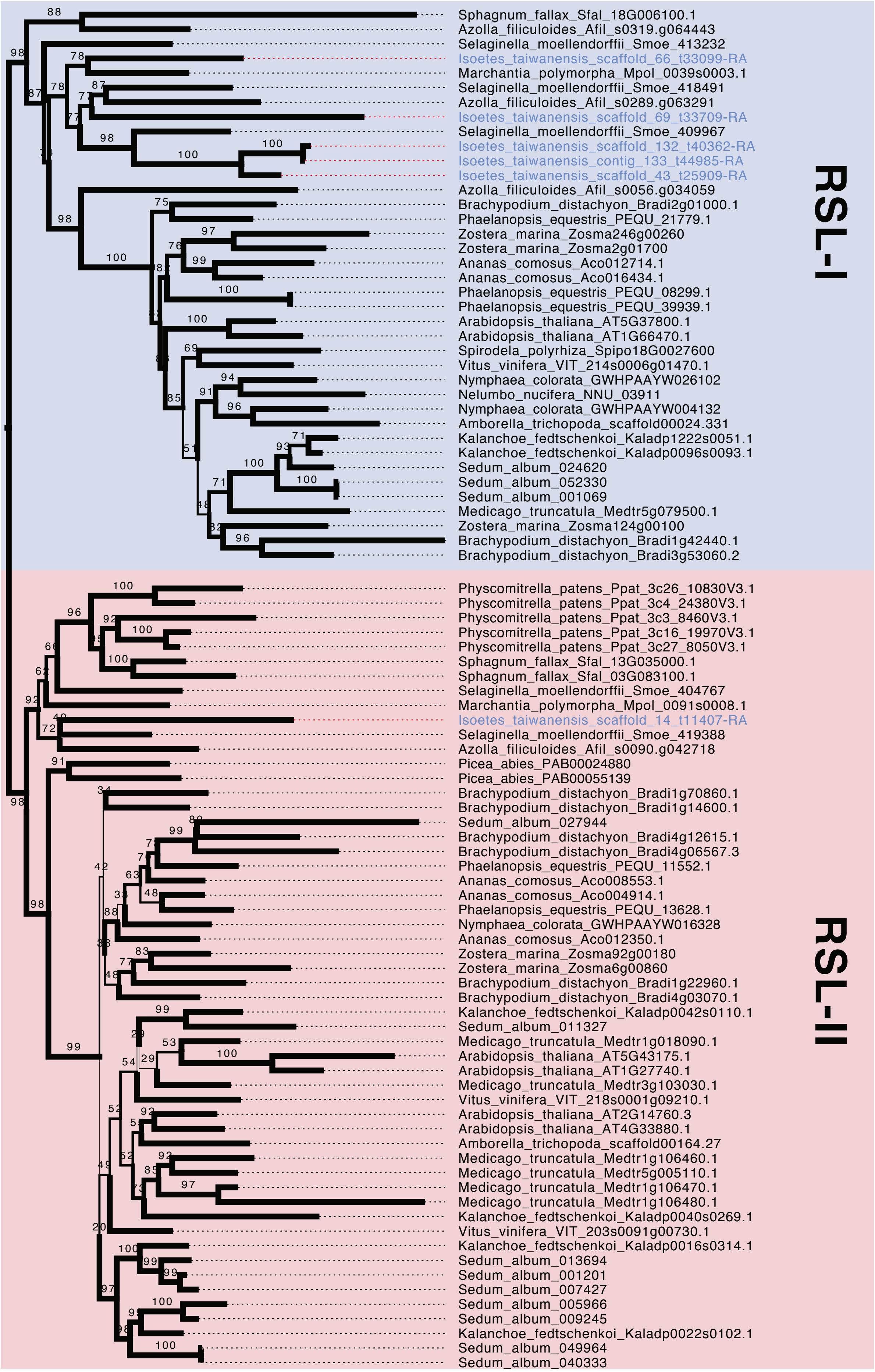


**Supplemental Figure S20:** ***ROOT HAIR DEFECTIVE SIX-LIKE*** **(*RSL*) genes are present in *Isoetes taiwanensis.*** A phylogeny showing the placement of Class-I and Class-II RSL genes found in *I. taiwanensis* relative to other land plants. Branch thickness corresponds to bootstrap support.

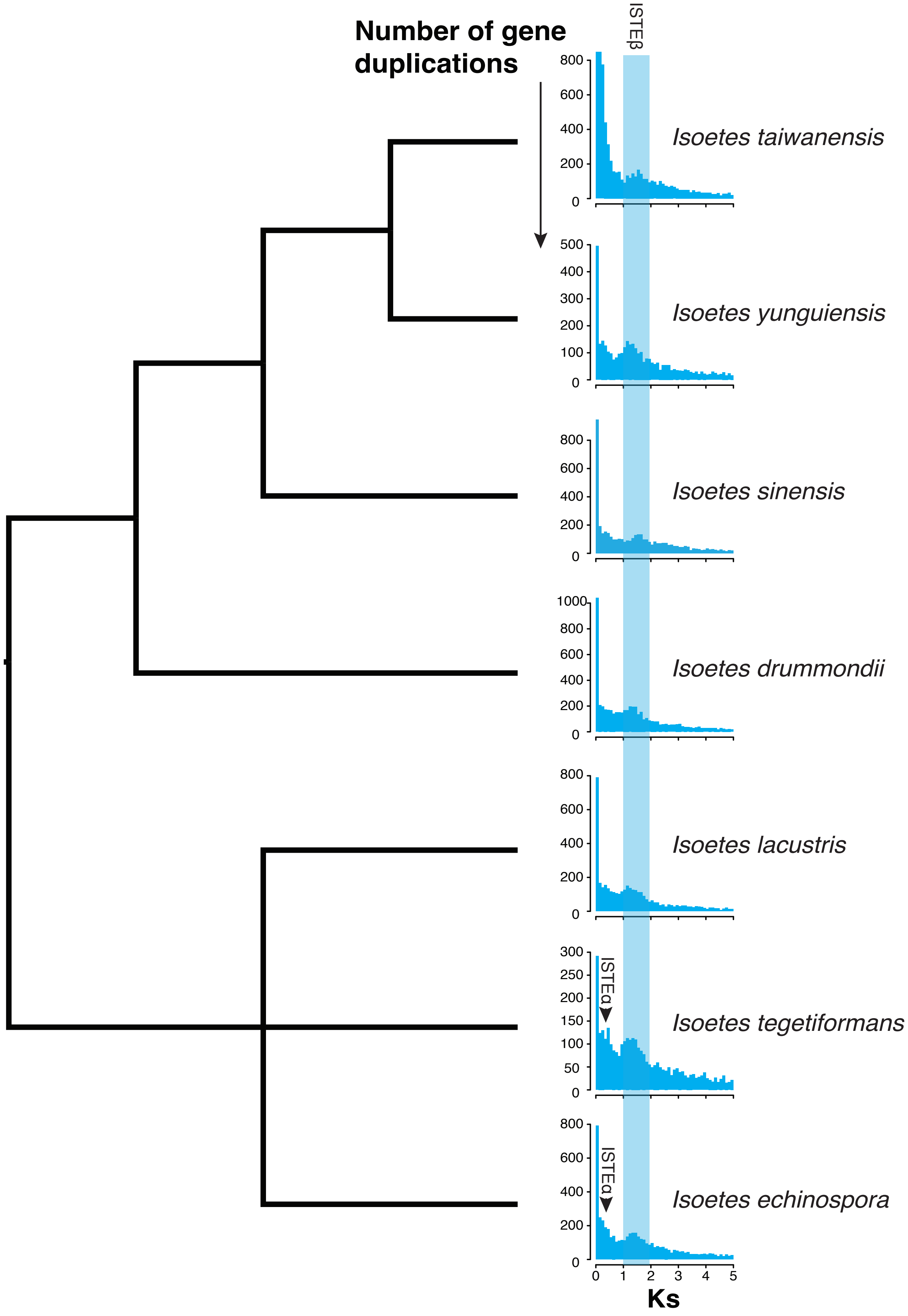


**Supplemental Figure S21: A whole genome duplication predates the divergence of two major *Isoetes* clades.** Multiple species of *Isoetes* exhibit a Ks peak near Ks = 1.5-1.8 (ISTEβ), indicated by the blue box. A Ks peak representing a proposed more recent gene duplication identified in *Isoetes tegetiformans* and *Isoetes echinospora* (ISTEα) is indicated by black arrows. The Ks plot for *I. taiwanensis* was generated from genomic data and the initial peak truncated to highlight ISTEβ peak. All other species were generated from transcriptomic data.

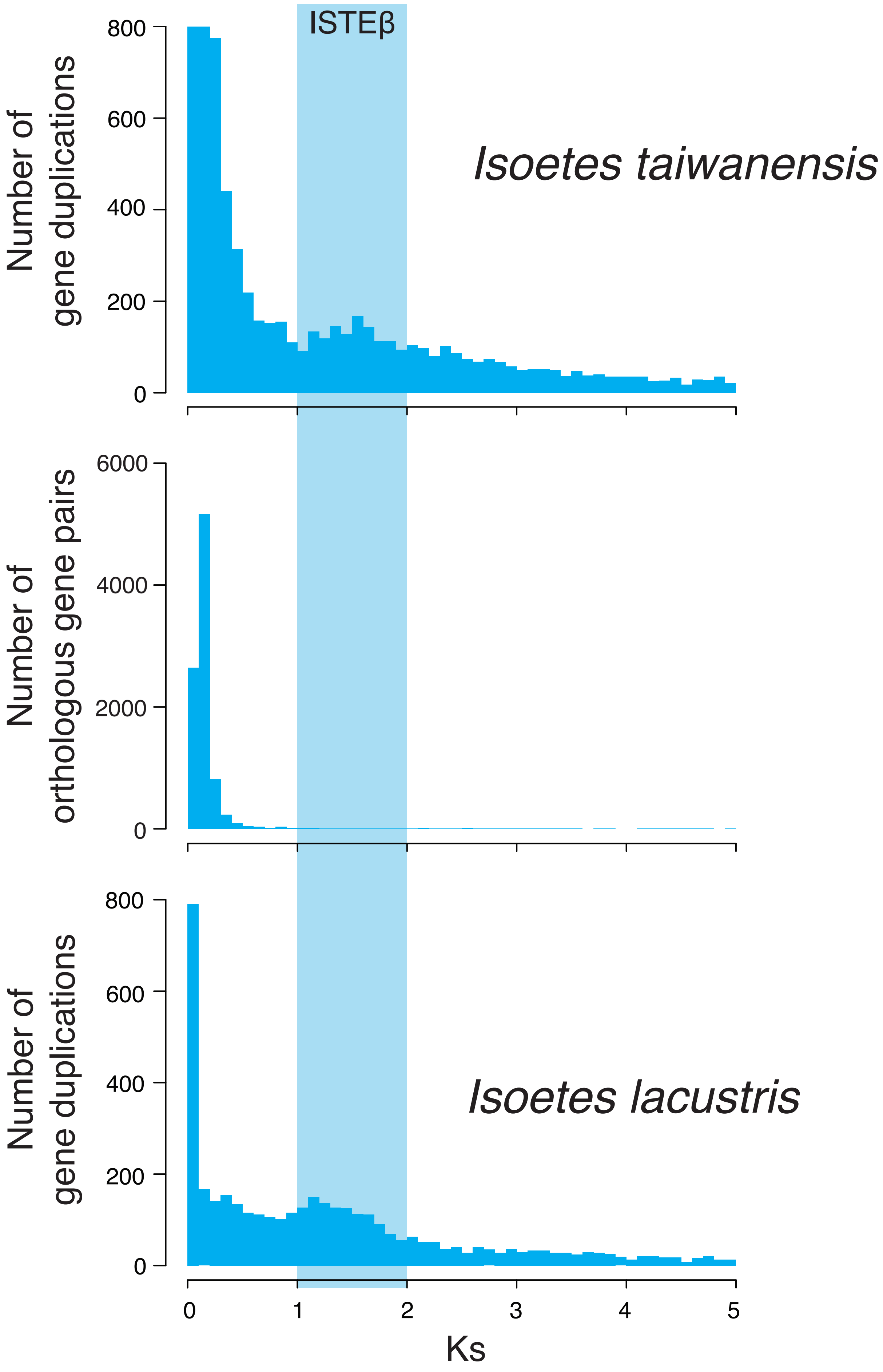


**Supplemental Figure S22: A whole genome duplication predates the divergence of two major *Isoetes* clades.** Comparison of ortholog divergence (Ks) between two distantly related species of *Isoetes* dates an ancient WGD (ISTEβ; blue box) to before the divergence of their respective clades. The Ks plot for *I. taiwanensis* was generated from genomic data and the initial peak truncated to highlight ISTEβ peak. The Ks plot for *I. lacustris* was generated from transcriptomic data.

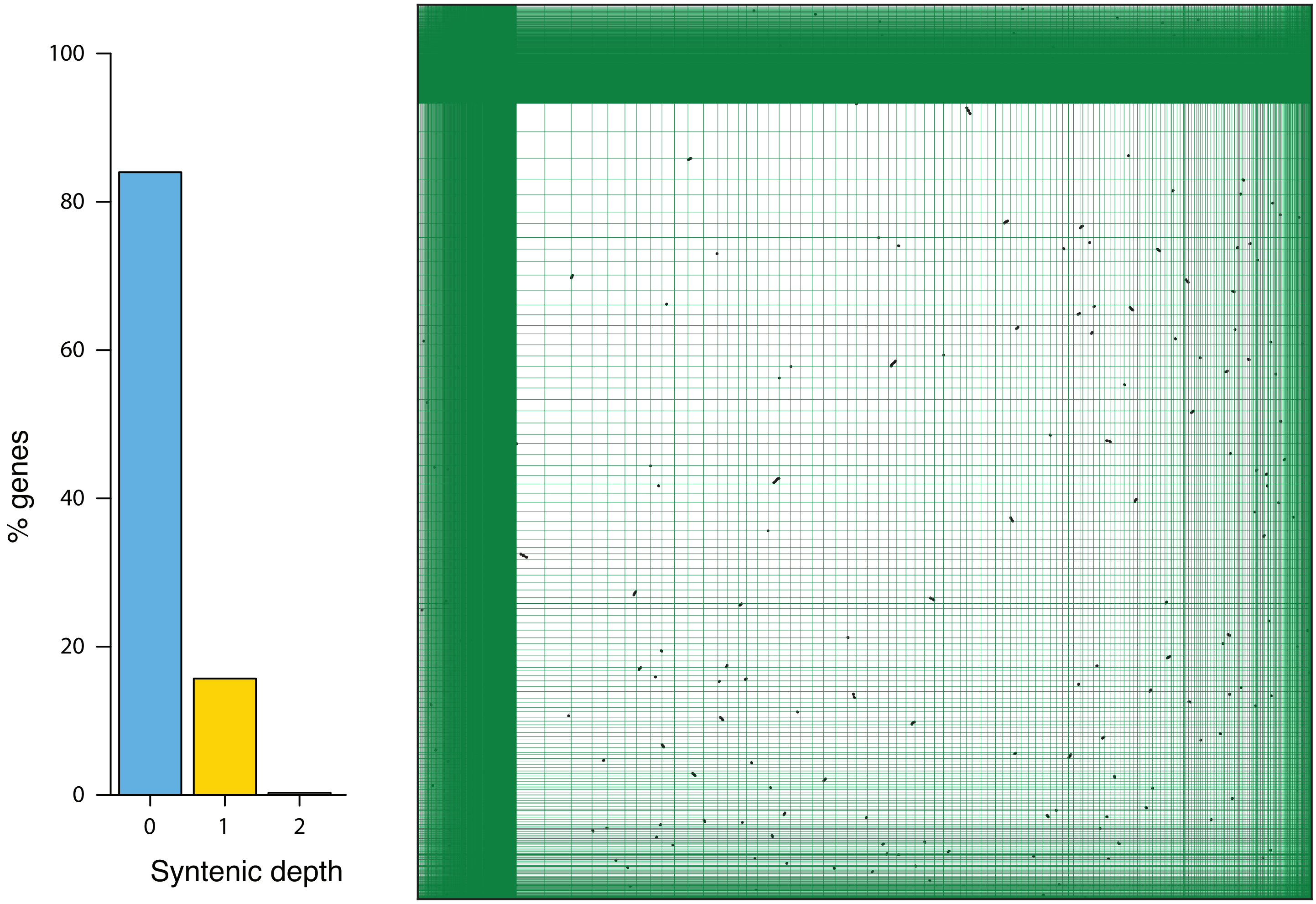


**Supplemental Figure S23: *Isoetes taiwanensis* self synteny plot.** Collinearity analysis identified 6,196 genes in 107 collinear blocks. The bar graph at the left shows the proportion of syntenic depth throughout the genome.

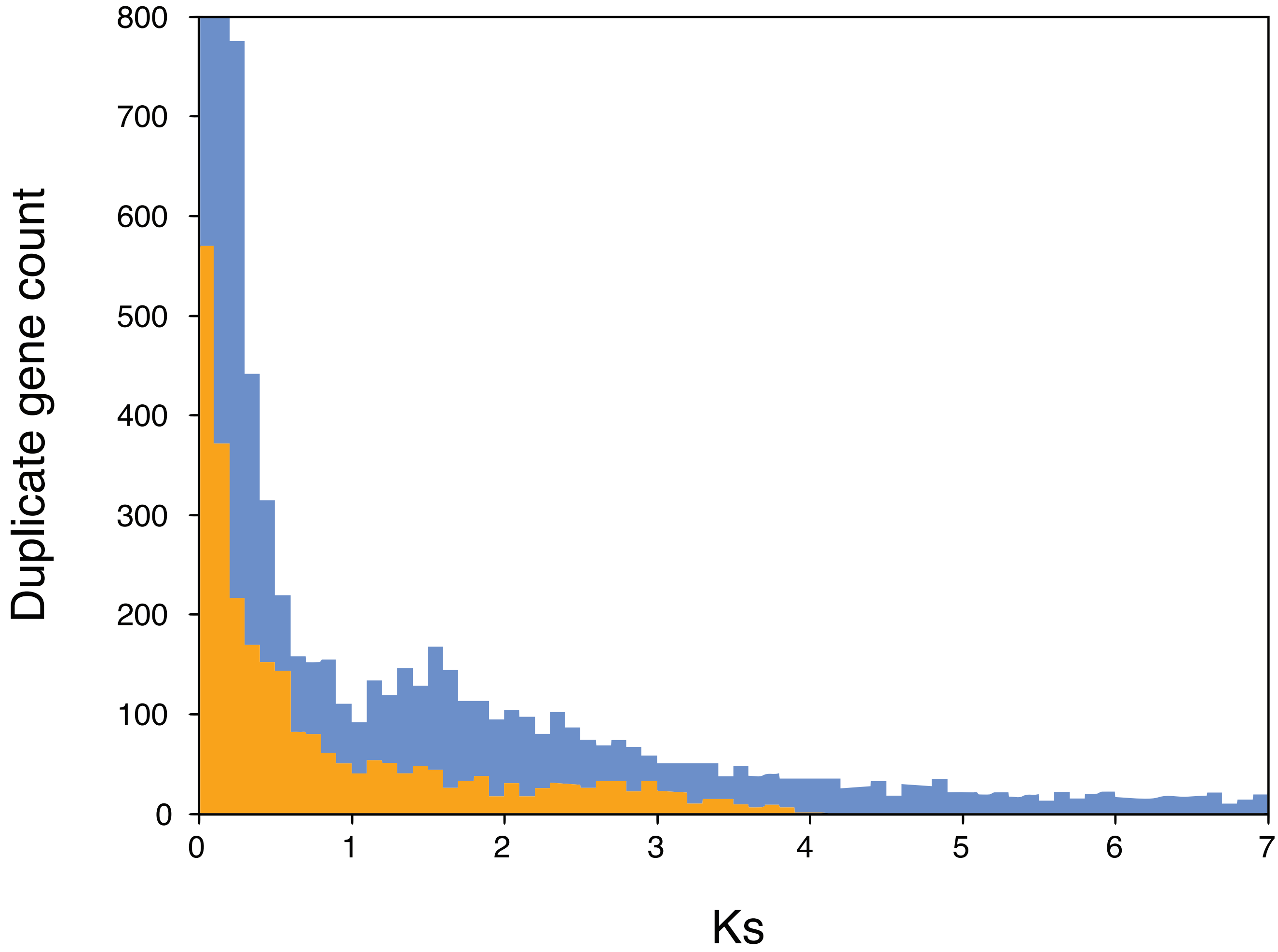


**Supplemental Figure S24: Ks distribution of syntenic gene pairs differs from that of the whole genome.** A peak in the genome-wide Ks distribution (blue) around 1.8 was not evident when Ks analysis was restricted to syntenic gene pairs (orange).

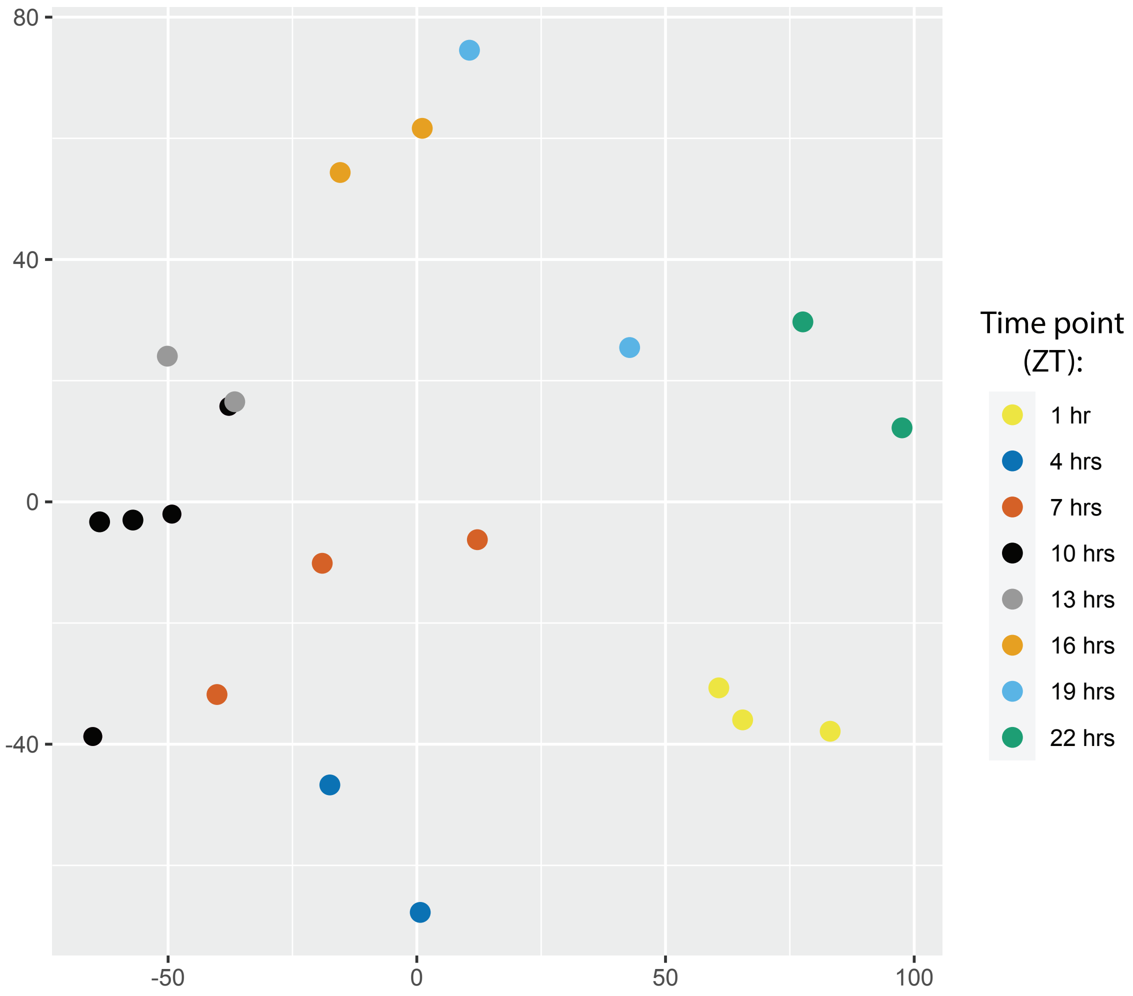


**Supplemental Figure S25: RNA-seq samples used for gene expression analyses cluster together by time point.** A Multidimensional scaling plot of RNA-seq data shows clustering by time point in a clockwise manner following the removal of outliers.

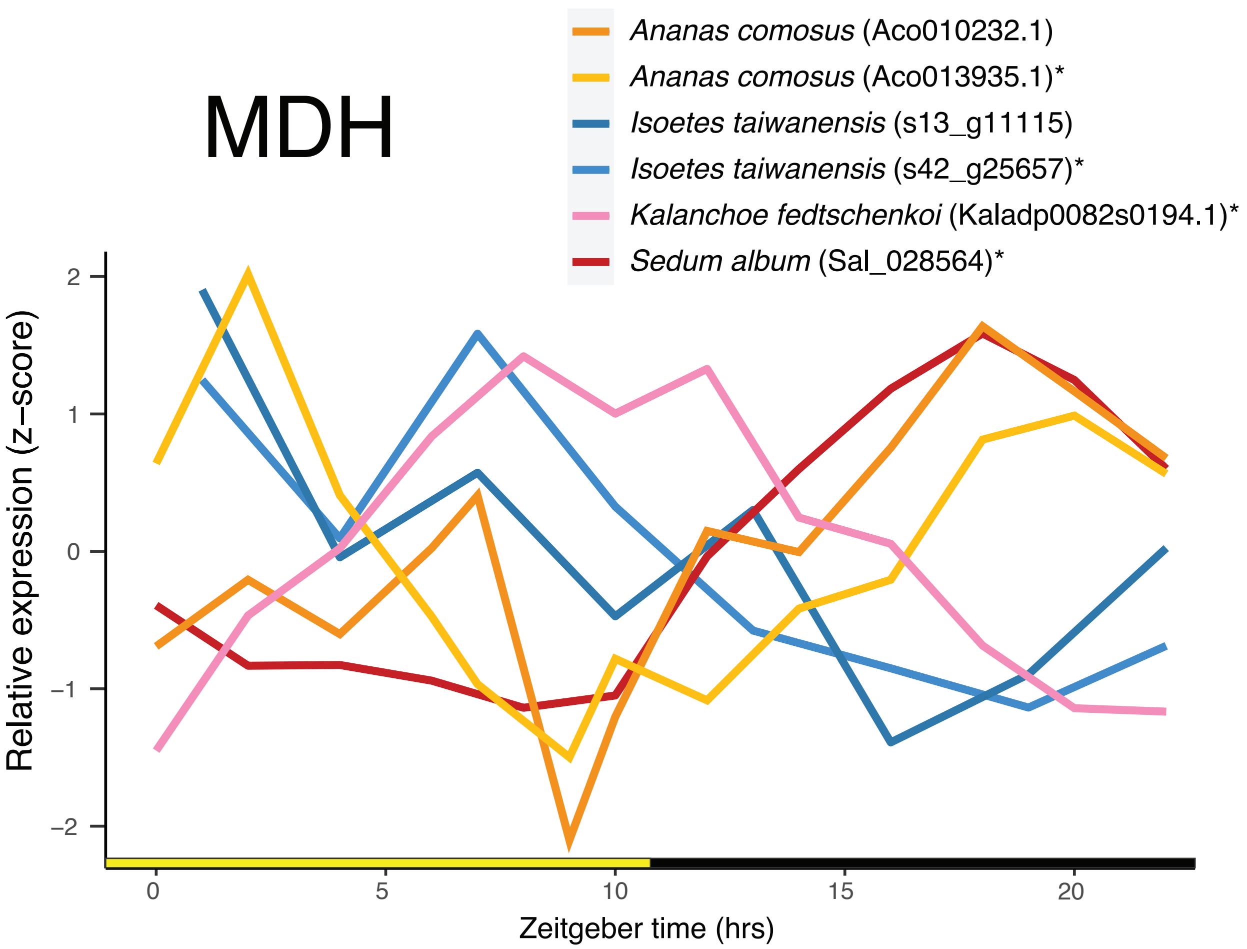


**Supplementary Figure S26: Comparison of cycling copies of MDH across CAM species.** Plot shows relative gene expression (z-score of TPM normalized RNAseq data) for all MDH copies that are known to cycle in terrestrial CAM plants as well as two copies from *Isoetes taiwanensis*. Orthologous genes are marked with an asterisk (*).

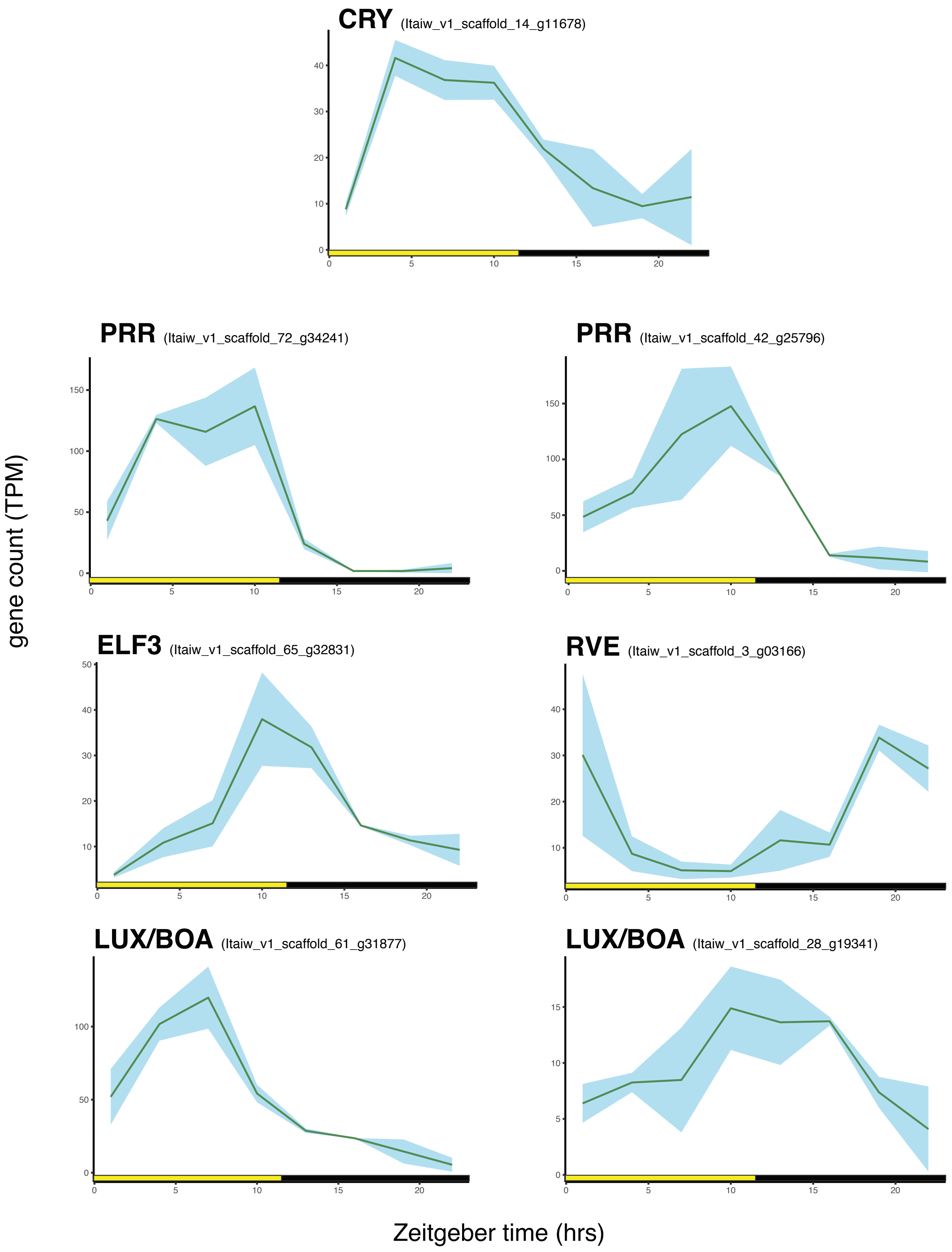


**Supplementary Figure S27: Many circadian associated genes exhibit typical cycling behavior in *Isoetes taiwanensis*.** Plots showing TOD expression of circadian associated genes in *I. taiwanensis*. Cycling genes are named for their nearest orthologues in Arabidopsis.
