## Supplementary Notes for "Underwater CAM photosynthesis elucidated by *Isoetes* genome"

#### *RNA editing in the Isoetes taiwanensis chloroplast genome*

The mitochondrial genome of *Isoetes engelmannii* was previously shown to have one of the highest numbers of RNA-editing sites known to date[1], in which a total of 1,782 sites were affected, and both the canonical C-to-U and the reverse U-to-C edits were found. However, it is unclear if the plastid genome of *Isoetes* also exhibits a similar level of RNA-editing. Here we mapped our RNA-seq data (that were not poly-A enriched) to the *I. taiwanensis* plastome, and identified a total of 465 RNA-editing sites. While 43 editing sites were located in noncoding regions, 422 were found in 70 protein coding genes. Of these, the majority were of the C-to-U type (399 sites) with U-to-C editing found at 23 sites. The total number of RNA-editing sites in *Isoetes* is comparable to plastomes of other seed-free plants, but substantially lower than *Selaginella*, where more than 3,400 sites were reported[2]. There appears to be a bias toward edits at the first codon position with 233, 56, and 122 editing events found at the first, second, and third positions respectively. Despite this, slightly more than half of edits (54%), including all identified U to C edits, were silent. The most heavily edited gene was *petN* with editing detected in 5.6% of its nucleotides. In addition, *psb* genes were generally highly modified, comprising 5 of the 10 most edited genes (Supplementary Figure S2). The reason behind such concentrated editing is unclear.

#### *Gene family evolution*

To assess gene homology between *Isoetes* and other plants, we conducted an Orthofinder analysis containing 25 species from across the plant phylogeny. Our analysis placed 647,535 genes into 40,144 orthogroups. Of those, 12,890 orthogroups contained at least one gene from *I. taiwanensis* with 3,391 being unique to *Isoetes*. A total of 6,241 *Isoetes* genes are singletons that were not assigned to any group. Surprisingly, *Isoetes* genes showed greater orthogroup overlap with seed plants than with any seed-free taxa (Supplementary Fig. S4). While this relationship may have been driven by the relatively high number of angiosperm taxa included in the original analysis relative to other groups, a similar trend was not seen in *Selaginella moellendorffii* (Supplementary Figure S4).

#### *Genes related to lignin biosynthesis*

One of the foremost features of vascular plants is lignification. The three major types of lignin monomers (hydroxyphenyl, guaiacyl, and syringyl; H-, G-, and S-lignin) are sourced from the phenylpropanoid pathway, and lycophytes use intriguing biochemical routes towards these compounds. In *Selaginella moellendorffii*, S-lignin is produced via two enzymes, CAFFEIC ACID/5-HYDROXYFERULIC ACID O-METHYLTRANSFERASE (SmCOMT) and FERULATE 5-HYDROXYLASE (SmF5H); both evolved independently from the canonical angiosperm COMT and F5H counterparts[3,4]. Similar to *S. moellendorffii*, *I. taiwanensis* lacks canonical *COMT* and *F5H* genes, but has orthologs of *SmCOMT* and *SmF5H* as well as the rest of lignin biosynthesis enzymes—offering putative routes towards all types of lignin (Supplemental Figures S5-S17). This result implies that the evolution of such an alternative S-lignin pathway might predate the divergence of *Selaginella* and *Isoetes*. However, the presence of S-lignin in *Isoetes* species appears ambiguous[5]. Residue analyses of *I. taiwanensis COMT* candidates found that, in contrast to the functionally characterized *SmCOMT* (Smoel_227279), the catalytic triad (HDE) contains a radical substitution (HNE) (Supplemental Figure S12); further, in contrast to all other gene families (Supplemental Figure S12 and S17), none of the *ItCOMT* homologs showed detectable gene expression. Future work on lignin biochemistry and enzyme assays are needed to clarify the evolution of S-lignin biosynthesis in lycophytes.

#### *Genes related to stomata development*

The presence of genes associated with stomatal development in *Isoetes* is of interest because the production of functional stomata varies throughout the genus. While some aquatic and at least one terrestrial species only produce astomatous leaves, some, including *I. taiwanensis*, do produce functional stomata under aerial conditions[6]. A recent study using transcriptomic data found evidence for loss of stomatal patterning genes *EPIDERMAL PATTERNING FACTOR 1/2* (*EPF1/2*) and *TOO MANY MOUTHS* (*TMM*)[7], even though this species is known to produce stomata[8]. In *I. taiwanensis*, while our genomic evidence corroborated the absence of *EPF1/2* orthologues, we did find two copies of *TMM*, one of which was expressed at low levels in leaf tissue (Supplementary Figure S18). In addition, we confirmed the presence of orthologues of other stomatal patterning genes, such as *SPEECHLESS/MUTE/FAMA* (*SMF*) and *FAMA* (Supplementary Figure S19). Thus, despite their aquatic growth habit, *Isoetes* appear to have retained much of the genetic machinery required for stomatal development (Supplementary Table S2). This is what we might expect given *Isoetes'* amphibious nature, and highlights the danger of inferring gene absence from transcriptomic data alone.

#### *Genes related to root development*

The homology of *Isoetes* roots has been the source of some controversy. *Isoetes* 'rootlets' are highly similar to the 'stigmarian' roots of ancient lycopsids and their superficial resemblance to aboveground structures has led some researchers to speculate that *Isoetes* 'rootlets' are in fact modified leaves[9]. However, a recent study[10] of the *I. echinospora* transcriptome identified *ROOT HAIR DEFECTIVE SIX-LIKE* (*RSL*) genes, which have been used as a marker for root development in vascular plants[11,12]. We found multiple copies of *RSL* genes in the *I. taiwanensis* genome and subsequent phylogenetic analysis placed 5 copies in the RSL Class I clade and a single copy in the RSL Class II clade (Supplementary Fig. S20). Taken together, our results are consistent with the earlier transcriptome-based study and provide evidence for homology of root structures across vascular plants, at least at the genetic level.

### References


1. Grewe, F. *et al.* A unique transcriptome: 1782 positions of RNA editing alter 1406 codon identities in mitochondrial mRNAs of the lycophyte Isoetes engelmannii. *Nucleic Acids Res.* **39**, 2890–2902 (2011).
2. Oldenkott, B., Yamaguchi, K., Tsuji-Tsukinoki, S., Knie, N. & Knoop, V. Chloroplast RNA editing going extreme: more than 3400 events of C-to-U editing in the chloroplast transcriptome of the lycophyte Selaginella uncinata. *RNA* **20**, 1499–1506 (2014).
3. Weng, J.-K., Li, X., Stout, J. & Chapple, C. Independent origins of syringyl lignin in vascular plants. *Proc. Natl. Acad. Sci. U. S. A.* **105**, 7887–7892 (2008).
4. Weng, J.-K., Akiyama, T., Ralph, J. & Chapple, C. Independent recruitment of an O-methyltransferase for syringyl lignin biosynthesis in *Selaginella moellendorffii*. *Plant Cell* **23**,

2708–2724 (2011).
5. Espiñeira, J. M. *et al.* Distribution of lignin monomers and the evolution of lignification among lower plants. *Plant Biol* **13**, 59–68 (2011).
6. Keeley, J. E. CAM photosynthesis in submerged aquatic plants. *Bot. Rev.* **64**, 121–175 (1998).
7. Harris, B. J., Harrison, C. J., Hetherington, A. M. & Williams, T. A. Phylogenomic Evidence for the Monophyly of Bryophytes and the Reductive Evolution of Stomata. *Curr. Biol.* **30**, 2001–2012.e2 (2020).
8. Keeley, J. E. Distribution of diurnal acid metabolism in the genus Isoetes. *Am. J. Bot.* **69**, 254–257 (1982).
9. Rothwell, G. W. & Erwin, D. M. The rhizomorph apex of paurodendron; Implications for homologies among the rooting organs of lycopsida. *Am. J. Bot.* **72**, 86–98 (1985).
10. Hetherington, A. J., Emms, D. M., Kelly, S. & Dolan, L. Gene expression data support the hypothesis that Isoetes rootlets are true roots and not modified leaves. *Sci. Rep.* **10**, 1–10 (2020).
11. Kim, C. M. & Dolan, L. ROOT HAIR DEFECTIVE SIX-LIKE Class I Genes Promote Root Hair Development in the Grass Brachypodium distachyon. *PLoS Genet.* **12**, e1006211 (2016).
12. Huang, L., Shi, X., Wang, W., Ryu, K. H. & Schiefelbein, J. Diversification of Root Hair Development Genes in Vascular Plants. *Plant Physiol.* **174**, 1697–1712 (2017).
